# Drainage of inflammatory macromolecules from brain to periphery targets the liver for macrophage infiltration

**DOI:** 10.1101/2020.06.18.159392

**Authors:** Linlin Yang, Jessica A. Jiménez, Alison M. Earley, Victoria Hamlin, Victoria Kwon, Cameron T. Dixon, Celia E. Shiau

## Abstract

Many brain pathologies are associated with liver damage, but a direct link has long remained elusive. Here, we establish a new paradigm for interrogating brain-periphery interactions by leveraging zebrafish for its unparalleled access to the intact whole animal for *in vivo* analysis in real time after triggering focal brain inflammation. We reveal that drainage of inflammatory macromolecules from the brain led to a strikingly robust peripheral infiltration of macrophages into the liver independent of Kupffer cells. We further demonstrate that this macrophage recruitment requires signaling from the cytokine IL-34, Toll-like receptor adaptor protein MyD88, and neutrophils. These results highlight the ability for circulation of brain-derived substances to serve as a rapid mode of communication from brain to the liver. Understanding how the brain engages the periphery at times of danger may offer new perspectives for detecting and treating brain pathologies.

## Introduction

Whether a diseased or injured brain transmits signals to the periphery to activate a response is an interesting prospect in understanding brain-periphery communication, but remains underexplored. Interestingly, liver damage and neutrophil recruitment to the liver are common features in sepsis-related injury[1] and several central nervous system (CNS) pathologies[2, 3], including traumatic brain injury, multiple sclerosis, and Alzheimer’s disease. A few recent studies have implicated a systemic, albeit most prominently a hepatic response to CNS inflammation due to CNS trauma or injury in mammals[4, 5]. However, the underlying mechanisms that may link brain inflammation with liver impairment remain unclear. The ability to directly track the cellular and molecular processes from the brain to the liver *in vivo* provides a direct means to understand the brain-liver association.

To investigate components of communication between the brain and periphery, we hypothesized that macrophages, key innate immune cells, capable of long-range migration and signaling[6], could act as mediators of brain-periphery communication. To this end, we investigated if a brain perturbation such as inflammation could trigger a peripheral organ response mediated by macrophages. We employed the zebrafish because it offers unparalleled access to *in vivo* tracking and manipulation of molecular and cellular processes in the intact whole vertebrate animal from brain to peripheral organs, which are largely conserved from zebrafish to human[7]. Using zebrafish, we were able to directly capture the dynamic changes in macrophages occurring in the body after focal brain challenge and found the liver to be the most prominent target peripheral organ for immune infiltration. Our data supports the notion that infiltrating and resident macrophages in the liver may critically modulate CNS and systemic inflammation by shaping the hepatic response to circulating or widely distributed molecules that may be infectious, toxic, or exogenous.

A possible route through which the brain may affect liver function may simply be a drainage of effector molecules into circulation, albeit even at a trace level, to initiate systemic inflammation, rather than direct brain to liver signaling. While much research has focused on mechanisms penetrating the blood-brain barrier (BBB) or the blood-cerebrospinal fluid (BCSF) barrier to enable entry of a peripheral agent into the brain parenchyma such as those relating to infectious diseases causing brain dysfunction (for example, by neurotropic viruses including *Rabies lyssavirus*, West Nile virus, and cytomegalovirus) and drug delivery to the brain [8, 9], far less attention has been given to investigating the reciprocal transfer from brain to circulation and its consequences. Limiting the free passage of solutes and large molecules into the CNS is tightly-regulated by both BBB and BCSF barriers to ensure protection of the CNS from inappropriate tissue damage and inflammation. By contrast, the removal of waste and toxic agents from the brain interstitial space to ensure normal brain health and function requires appropriate metabolism or efflux of these substances [10]. Previous studies on movement of CNS fluid and solutes indicate several possible routes of drainage of substances from the brain parenchyma including perivascular pathways that may exit through the CSF or lymph tracts, and the BBB[10]. Therefore, if the net outcome is some degree of systemic inflammation due to an efflux of inflammatory cues from the brain to circulation, even at a low level, then it could effectively target the liver, but this remains to be investigated.

Given that the liver is equipped to process a large fraction of the total blood circulation from two major blood supplies[11], the hepatic artery and portal vein from the gastrointestinal tract, it may not be surprising that the liver would be highly sensitive and responsive to inflammatory mediators and foreign agents in the blood flow. In fact, common and infectious bacteria (including *Escherichia coli, Klebsiella pneumoniae, Salmonella typhimurium* and *Listeria monocytogenes*) in the bloodstream have often been found to be cleared by the liver [12, 13]. However, the mechanisms that specifically make the liver susceptible to systemic inflammation remain incompletely understood. Besides metabolic functions, the liver provides critical immune surveillance by recognizing and clearing away infectious, toxic, and microbial substances in the blood, a function that has largely been attributed to the Kupffer cells [14-16]. Systemic inflammation stemming from infection, toxic insults, and autoimmunity [17] can cause chronic infiltration of leukocytes into the liver leading to liver damage and subsequent progression to fibrosis, cirrhosis or liver cancer [18-20]. However, how the liver responds and contributes to systemic inflammation by way of leukocyte infiltration remains poorly understood, a process that has not been directly visualized *in vivo* for an open dissection.

Here, we reveal new insights into the cellular dynamics and critical roles of the IL-34 and MyD88 signaling pathways as well as Kupffer-cell-independent mechanisms in mediating immune infiltration of the liver in response to systemic lipopolysaccharides (LPS), classic pro-inflammatory bacteria-derived stimuli, after brain intraparenchymal LPS microinjection. We show that inflammatory cues could originate from the brain and trigger immune infiltration of the liver, a process involving drainage of molecules from brain to circulation not previously appreciated. Using comparative analyses, time-lapse imaging, and blocking circulation, we found that the effects of brain LPS injection were largely recapitulated by intravenous LPS injection, and stemmed from systemic inflammation akin to sepsis or endotoxemia. We can block infiltration of immune cells into the liver by disrupting MyD88, a key adaptor of Toll-like receptors responsible for LPS recognition, or by eliminating the cytokine IL-34 pathway, and by reducing inflammation pharmacologically. Additionally, coordination between macrophages and neutrophils is also essential for the liver infiltration. Infiltration by macrophages and neutrophils negatively impacts the liver by promoting inflammation, and disrupting normal hepatic growth. Taken together, changes in macrophage behavior involving immune cell infiltration of the liver may be triggered by drainage of inflammatory cues from the brain, providing a possible readout for an altered brain.

## Results

### Brain immune activation is associated with macrophages infiltrating the liver prior to Kupffer cell establishment

Reciprocal connections between brain and liver are apparent in various conditions, including encephalopathy and encephalitis after severe liver damage[21, 22], liver disruption after traumatic brain injury[23, 24], and intracerebral injection of pro-inflammatory cytokines[4]. However, the routes of communication directly linking brain to liver remain poorly understood. To investigate one possible avenue of this, we sought to determine whether a brain perturbation such as inflammation could trigger macrophage activities corresponding to a response by the liver or other peripheral organs. To this end, we directly microinjected bacterial lipopolysaccharides (LPS) or *E. Coli* cells, well-established immune activators, into the brain tectum at 4 days post-fertilization (dpf) at a stage known to have a well formed brain with an established BBB [25] and choroid plexus/ventricle system [26, 27] as previously described[28]. Stage-matched animals that were not injected or injected with the water vehicle in the brain were used as the control group (Figure 1 and Supplementary Fig. 1). To analyze a possible change in peripheral macrophages after brain-LPS injection at 4 dpf, we used whole-mount RNA *in situ* hybridization for a macrophage marker, *mfap4*, to characterize the macrophage distribution in the whole body. Strikingly, we found a robust and distinctive aggregation of macrophages in the liver and near the brain injection site, but not apparent elsewhere (Supplementary Fig. 1). By contrast, the uninjected and water injected controls at 4 dpf were devoid of macrophages in the liver (Supplementary Fig. 1). We also found at least some of these macrophages in the liver to be activated based on their expression of the mitochondrial enzyme gene *irg1/acod1* (Supplementary Fig. 1) known to be highly upregulated during inflammation and specifically induced in inflammatory macrophages in zebrafish[29, 30]. Both injections of LPS or live *E. coli* cells into the brain led to macrophage presence in the liver, albeit LPS effects were consistently stronger (Supplementary Fig. 1).

**Figure 1.**
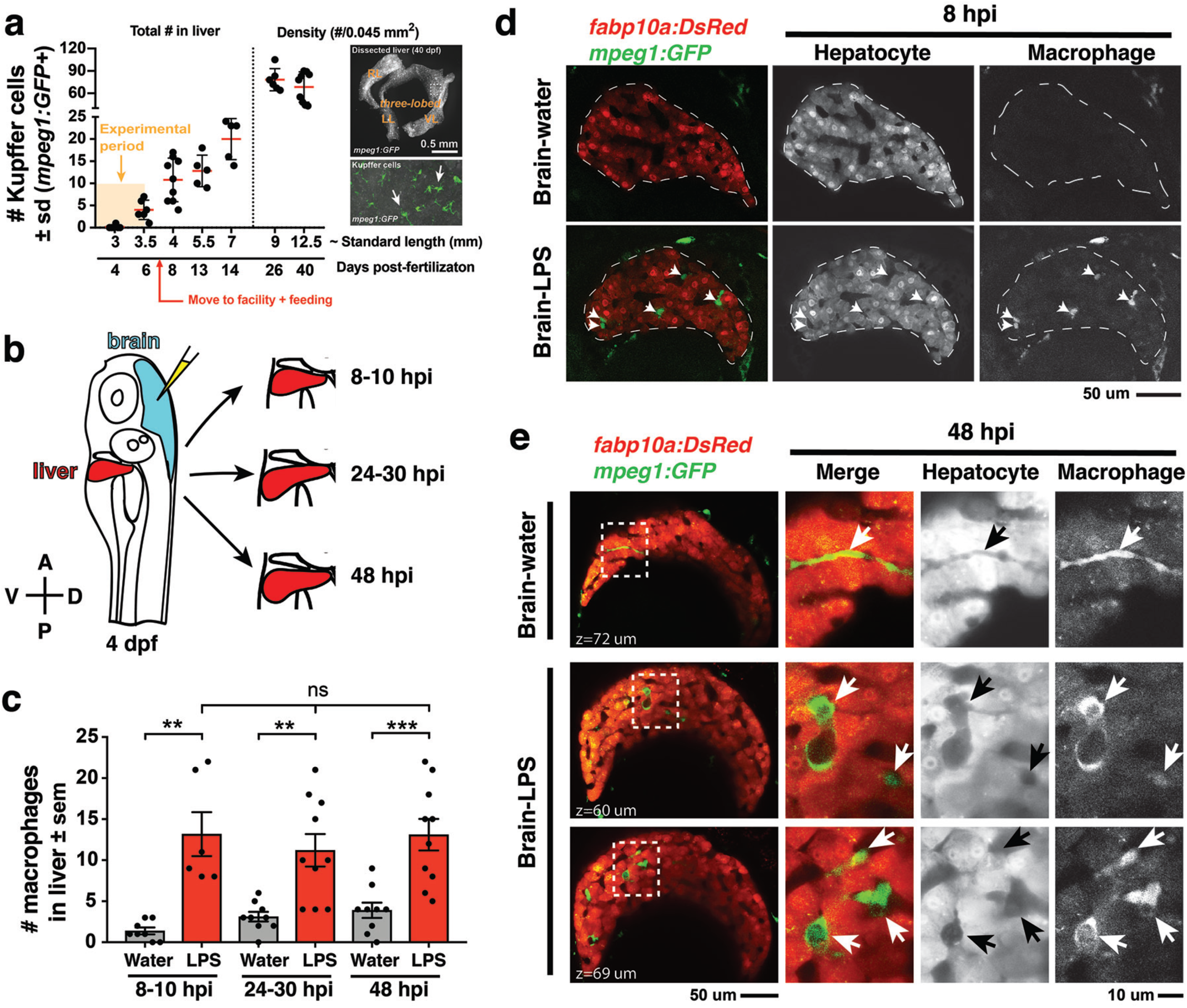
Induction of brain inflammation triggers macrophage infiltration into the liver. **a** Time-course of Kupffer cell development. Total macrophage numbers (*mpeg1:GFP*+) per liver from 4 to 14 days post-fertilization (dpf) and macrophage density (number per field of view) from dissected liver at juvenile adult stages from 26 to 40 dpf. Standard length corresponding to the stages shown. Kupffer cells are not present at 4 dpf at the time of brain microinjection and during most of our experimental period (orange box). Feeding began after 6 dpf to ensure normal growth. Fluorescent images on the right show dissected whole liver with the typical three-lobed structure at 40 dpf (top) and high magnification of top dotted box region showing Kupffer cells (bottom). LL, Left lobe; RL, right lobe; VL, ventral lobe. **b** Schematic of brain microinjection at 4 dpf and analysis of the hepatic response at 8-10 hours post injection (hpi), 24-30 hpi, and 48 hpi. A, anterior; P, posterior; V, ventral; D, dorsal. **c** Quantification of macrophage infiltration in the liver comparing between LPS and control water injections in the brain at different timepoints. **d** At 8hpi, single-plane image from a z-stack shows infiltrated macrophages (GFP+, arrows) nested between hepatocytes (DsRed+) in the liver (dotted region) after brain-LPS injection, but no macrophages observed in control brain-water injection. **e** At 48 hpi, images from two separate z-planes show a large number of infiltrating macrophages persist after brain-LPS injection (arrows), while few presumably Kupffer cells begin to appear in control brain-water injected animals at this late timepoint (arrow). Two-tailed Welch’s t-test was used to determine statistical significance for each pair-wise comparison. One-way ANOVA test for comparing the three LPS injection groups. sem, standard error of means; ns, not significant; **, p<0.01; ***, p<0.001.

To relate macrophage presence in the liver after brain-LPS injection to macrophages that normally reside in the liver (which we refer to as Kupffer cells), we analyzed the developmental timing of Kupffer cells, which had not been previously described, starting at the 4 dpf larval stage to the 40 dpf juvenile adult stage (Fig. 1a). We found normally an absence of Kupffer cells at the stage of our brain microinjection at 4 dpf (0.2 ± 0.4 standard deviation (s.d.) per liver), few to none at 5-6 dpf (4.0 ± 2.2 s.d. per liver), and once larvae were fed and moved into the fish facility, Kupffer cell numbers grew substantially (78.2 ± 14.9 s.d. per area of liver at 26 dpf) reaching to hundreds per liver in the juvenile adults (26-40 dpf at 9-12.5 mm standard length)(Fig. 1a). Prior to this work, Kupffer cells were thought to be missing or sparse in zebrafish and other teleost species [31, 32], but recent work tracing adult zebrafish Kupffer cells to their hematopoietic origin[33], and the data presented here collectively provide the first evidence for the prevalence of Kupffer cells in the zebrafish liver akin to their mammalian counterpart. Since the brain-LPS injection and subsequent analysis were conducted at 4 dpf, before the establishment of Kupffer cells, the presence of macrophages in the liver was likely because of active recruitment of peripheral macrophages and monocytes.

We subsequently conducted a time-course analysis of macrophage activities after brain-LPS injection at 4 dpf to determine if macrophages were actively infiltrating the liver. Using *in vivo* static and time-lapse imaging in double transgenic zebrafish expressing both the macrophage reporter *mpeg1:GFP* and the liver hepatocyte reporter *fabp10a:DsRed*, we observed macrophages actively migrating or circulating into the liver, affirming the *in situ* results. We captured macrophage dynamics in the liver region at 8-10 hours post-injection (hpi) of LPS in the brain (Supplementary Movie 1 and Supplementary Fig. 2) when substantial numbers of infiltrating macrophages can be observed. Conversely, the brain-water injected controls mostly had zero macrophages in the liver (average of 1.4 ± 1.2 standard deviation (s.d.) macrophages per liver) compared with an average of 13.2 ± 6.6 s.d. macrophages per liver after brain-LPS injection (Fig. 1 b-e). Later at 24-30 hpi and 48 hpi, significant numbers of macrophages in the liver persisted even two days after brain-LPS injection (11.2 ± 6.3 s.d. and 13.1 ± 6.1 s.d. macrophages per liver, respectively) (Fig. 1c, Supplementary Movie 2 and Supplementary Fig. 2). At these later timepoints, a few Kupffer cells may begin to emerge at less than 5 per liver (Fig. 1c). In agreement with the timeline of Kupffer cell development (Fig. 1a), only a few macrophages were detectable in uninjected and water-injected controls at the two later timepoints (3.1 ± 1.8 s.d. and 3.9 ± 2.8 s.d. macrophages per liver, respectively)(Fig. 1c). Macrophages in the liver at 48 hpi appeared more stationary than earlier at 8-10 hpi (Supplementary Movies 1 and 2, and Supplementary Fig. 2). In all timepoints of analysis, infiltrated macrophages in the liver were found in several locations, including inside the sinusoids (liver microvessels) similar to that described during mammalian liver injury[34], and surprisingly also in the parenchyma intermingling with hepatocytes (Fig. 1d-e, Supplementary Movies 1 and 2, Supplementary Fig. 2), a macrophage behavior previously not known. Due to some examples of broad liver *mfap4 in situ* expression (Supplementary Fig. 1) in comparison to a discrete number of infiltrating macrophages by live imaging after brain-LPS injection (Fig. 1), we assessed whether this could be explained by an induction of ectopic *mfap4* expression in the liver upon LPS activation (Supplementary Fig. 3). Using a transgenic line *mfap4:tdTomato* to mark cells expressing the *mfap4* gene, we found *mfap4* restricted to macrophages and absent in liver cells (Supplementary Fig. 3), suggesting the broad liver *mfap4* expression may be diffuse *in situ* signals coming from a liver more densely populated by infiltrated macrophages. To analyze the nature of physical contact of infiltrated macrophages to the hepatic sinusoids after brain-LPS injection, we imaged double transgenic zebrafish expressing the endothelial (*kdrl:mCherry*) and macrophage (*mpeg1:GFP*) reporters. Time-lapse imaging showed that macrophages can actively infiltrate the hepatic sinusoids, as well as be associated or entirely independent of the hepatic vasculature (Supplementary Fig. 4 and Supplementary Movie 3).

To determine whether this infiltration endured over time, we examined the number of liver-infiltrating macrophages over a two-day period after brain-LPS injection. Results indicate no significant change in macrophage presence in liver, suggesting that infiltrated macrophages either stayed in the liver, moved in and out of the liver at similar rates, or both. To distinguish these possibilities, we tracked these infiltrating macrophages directly *in vivo* over a continuous 10-hour period after brain-LPS injection at 4 dpf starting at 12 hpi. Using live cell tracking, we found a small number (∼13%) of infiltrating macrophages that stayed in the liver longer than 2 hours, including occasional infiltrates which remained in the liver past the total duration of imaging (> 9 hours) (Supplementary Fig. 5 and Supplementary Movie 4). Infiltrating macrophages were coined “residing” when they occupied the liver for more than 2 hours, and these were on average moving slower at 1.1 ± 0.7 um/min than the “transient” population at 2.2 ± 1.1 um/min which occupied the liver for more than one timepoint (2 minutes) but less than 2 hours (Supplementary Fig. 5). We found a significant fraction (∼30%) of infiltrating macrophages to be of the “transient” type, while the majority at 57.4% were circulating (detected only in a single timepoint)(Supplementary Fig. 5). Both the “residing” and “transient” populations were not truly stationary but rather moved dynamically back and forth across the liver parenchyma amounting to large total distances traveled over time (62.8 ± 66.7 um s.d. and 309.2 ± 285.9 um s.d. in 560 minutes of tracking)(Supplementary Fig. 5 and Supplementary Movie 4). These results indicated that both the presence of short- and long-term occupying macrophages accounted for the sustained large macrophage number in the liver even two days after brain-LPS injection (Fig. 1c).

### Microinjection of LPS into brain leads to systemic LPS distribution triggering immune infiltration of the liver

To understand how LPS in the brain may lead to liver effects, we first determined whether the injected LPS remained restricted to the brain or possibly transferred to the periphery over time. We used fluorescently tagged LPS to directly track the LPS molecules after brain tectum microinjection in the whole body for a continuous 24-hour period (Fig. 2 and Supplementary Movie 5). This provides a means to visualize binding, transport, and internalization of LPS in the brain and body. Fluorescently tagged dextran was used as a control tracer to analyze the general molecular distribution independent of LPS (Fig. 2 and Supplementary Movie 6) and to verify successful injections. Initially the LPS macromolecules injected into the brain parenchyma were restricted to the focal location of the injection site but quickly within seconds they filled the cerebral ventricles joining the cerebrospinal fluid (CSF) as they continue to flow into the spinal canal in an anteroposterior direction (Fig. 2 and Supplementary Movie 7), thus our zebrafish brain microinjections are comparable to mammalian intracerebroventricular injections[35].

**Figure 2.**
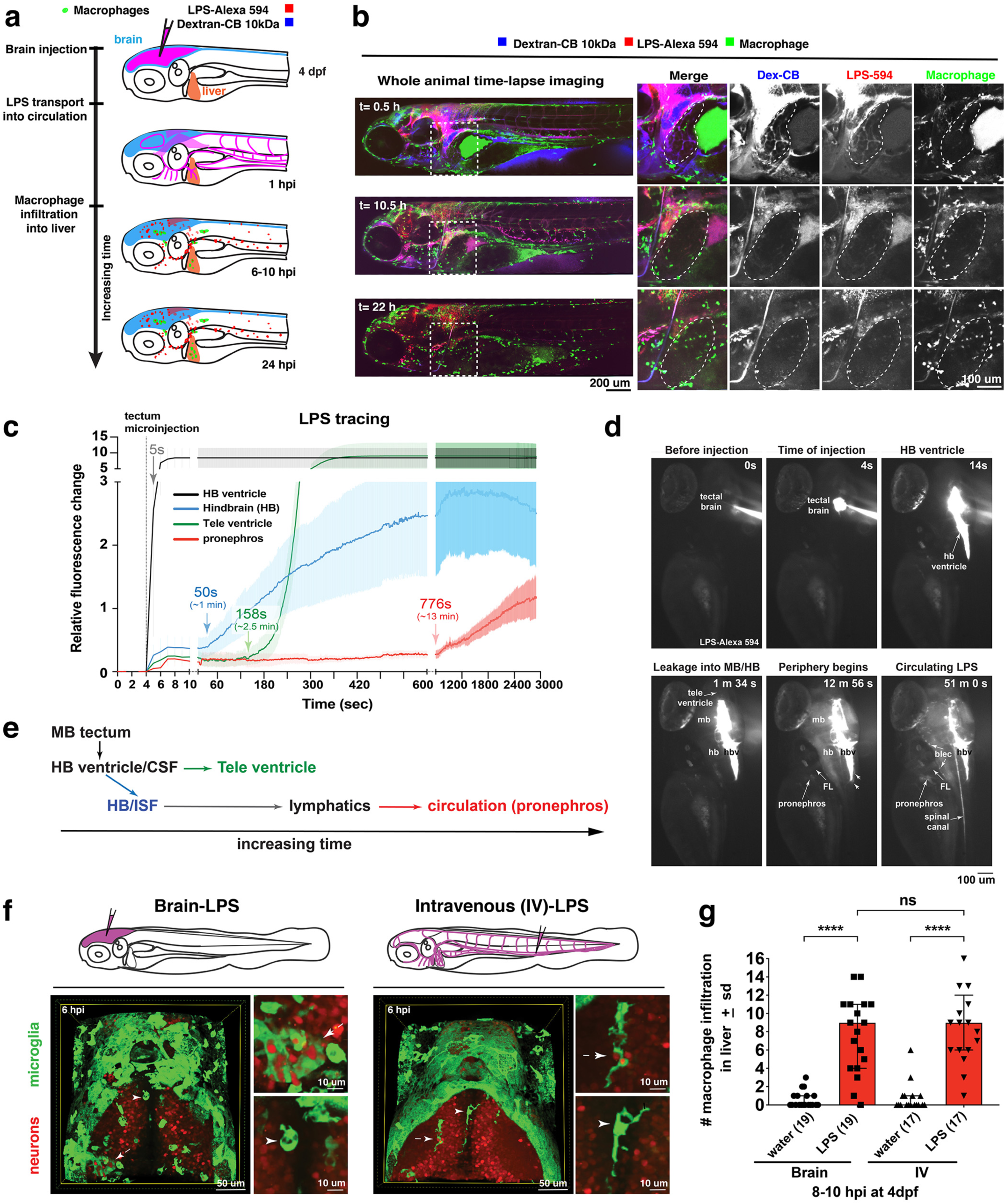
Brain LPS microinjection leads to drainage of LPS molecules into circulation, and causes a hepatic response similar to intravenous LPS injection. **a** Schematic showing time-course of LPS and macrophage distribution. Fluorescently tagged LPS-Alexa 594 shown in red, co-tracer dextran-cascade blue shown in blue, and detection of both is shown in magenta. **b** Left, representative 3D images from three timepoints of a 24-hour time-lapse imaging of a large frame stitched from four z-stack tiles (corresponding to Supplementary Movie 5). Right, high magnification of the 3D images in dotted box region on the left panel showing liver (dotted region) and surrounding area. Merged overlays and individual channels showing Dex-CB (blue), LPS-594 (red), and macrophages (*mpeg1:GFP*+). **c** Live recording of the brain microinjection at 4 dpf using Alexa 594 or Alexa 488 conjugated LPS was conducted to trace the distribution of LPS in real time at 1 frame per second using an automated acquisition software on a Leica M165 FC stereomicroscope with a high speed and high sensitivity deep-cooled sCMOS camera (DFC9000 GT). Kinetic time plot of relative fluorescence change ± sem of fluorescently tagged LPS starting at 0 seconds before the injection. Time of injection was at the 4 seconds timepoint. Arrows indicate the timepoint at which initial LPS signals were detected in the corresponding anatomical location. In some injected animals, LPS also flowed anteriorly from the midbrain ventricle into the telencephalon ventricle starting at about 2.5 minutes after injection. Measurements were conducted in three independent injected animals. **d** Still images representing key events of the dispersion of LPS starting from before to nearly 1 hour after the brain microinjection corresponding to Supplementary Movie 7. hb, hindbrain; tele, telencephalon; hbv, hindbrain ventricle; mb, midbrain; FL, facial lymphatics; blec, brain lymphatic endothelial cells; CSF, cerebrospinal fluid; ISF, interstitial fluid. **e** Schematic showing the major route through which LPS were transferred from the site of brain microinjection to peripheral circulation. **f** Top, illustration of brain and intravenous LPS injections. Bottom, 3D tectum brain volume from confocal live imaging at 4 dpf at 6 hours after brain or intravenous LPS injection using a 40x objective. Microglia (*mpeg1:GFP*+) and surrounding neurons (*nbt:DsRed*+) shown. Small panels show high magnification of microglia (arrows) and neurons corresponding to arrows in the large 3D brain volume image on the left. LPS injection in the brain led to a striking morphological activation of rounded and clustering microglia (arrows), but not by intravenous injection of LPS at 6 hpi. Superficial planes of the head are eliminated to better visualize the internal microglia, because the cranial skin surface is highly auto-fluorescent in the GFP channel. **g** Quantification of macrophage infiltration at 8-10 hpi in the 4 dpf zebrafish larvae. Two-tailed Welch’s t-test was used to determine statistical significance. sd, standard deviation; ns, not significant; ****, p<0.0001; LPS-594, LPS-Alexa 594; Dex-CB, cascade blue conjugated dextran, sem, standard error of means.

To evaluate the dynamics and major routes of LPS passage to the general circulation, we used high-speed and high-sensitivity stereomicroscopy to trace the movement of fluorescently tagged LPS in real time at 1 frame per second starting before the brain tectum microinjection to almost one hour after the injection (Fig. 2c-d and Supplementary Movie 7). While most of the LPS remained restricted within the ventricular system, we found the hindbrain ventricle (hbv) to be the key region from which LPS spread into the parenchyma and surrounding interstitial space, especially in the dorsal-most portion of the junction between the hindbrain and spinal cord encompassing the dorsal longitudinal anastomotic vessels (DLAVs) and the dorsal longitudinal lymphatic vessel (DLLV) (Supplementary Figs. 6 and 7, and Supplementary Movie 7). LPS at a low level were detected to exude out from the hbv into the hindbrain interstitial fluid (ISF) at around 1 minute after injection (Supplementary Movie 7 and Fig. 2c) and reached a maximum of about 30% of LPS injection level at 50 minutes after injection (Supplementary Fig. 8). After a delay of about 13 minutes after injection, the first LPS fluorescence signals were faintly measured in the pronephros, a region assessed as a proxy for general circulation (Fig. 2c-d and Supplementary Movie 7). Peripheral LPS level increased over time but remained low at less than 15% of the LPS injection level (Supplementary Fig. 8).

**Figure 3.**
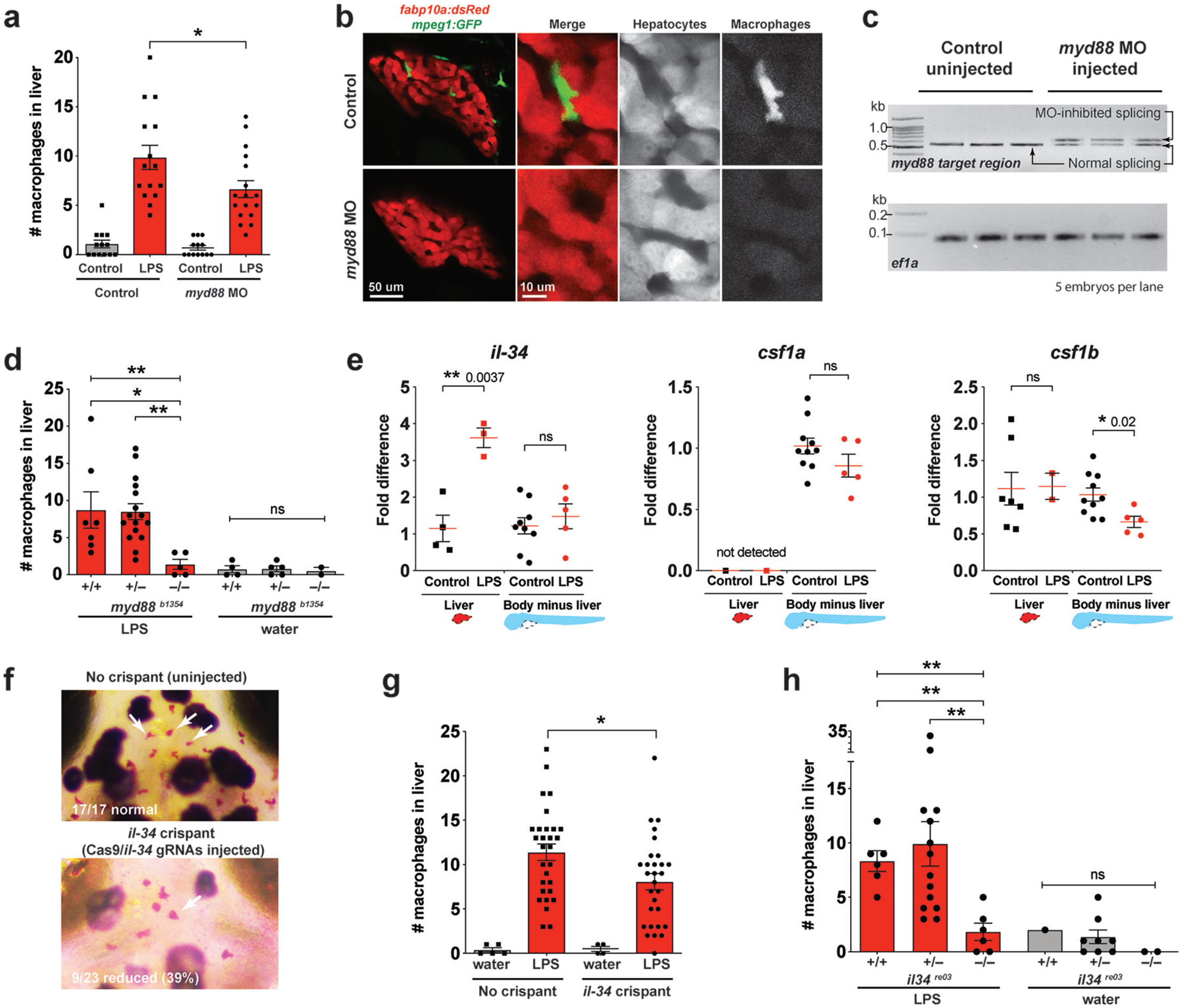
Infiltration of liver by macrophages triggered by brain perturbation is dependent on adaptor protein *myd88* and cytokine *il-34*. **a** Quantification of macrophage infiltration 6 hours after brain-LPS injection or control treatment (brain-water injection and no injection combined) in *myd88*-deficient morphants compared with control wildtype animals at 4 dpf. **b** Left column, representative single plane of whole liver (DsRed+) showing macrophage infiltration (GFP+) in the control animal but not in the *myd88* morphant. Second to fourth columns, high magnification of the merged overlay and single channels showing a single macrophage (GFP+) stationed between hepatocytes (DsRed+) in control but not in *myd88* morphant. **c** RT-PCR analysis showing efficacy of *myd88* morpholino in blocking normal *myd88* splicing at 3 dpf. Elongation factor 1 alpha (*ef1a*) PCR used as a sample quality control. **d** Complementary experiments using *myd88* genetic mutants derived from a heterozygous incross show either few or no macrophages in the liver after brain-LPS injection at 16-24 hpi similar to baseline brain-water injected animals, demonstrating a much stronger effect in reversing macrophage infiltration than the partial *myd88* knockdown by morpholinos. **e** qPCR analysis of *il-34, csf1a*, and *csf1b* expression in liver only and body-minus-liver tissues comparing brain-LPS injected animals with the control group (brain-water injected and uninjected animals combined) at 6 hpi in 4 dpf zebrafish. **f** Representative images of non-crispant (top) and *il-34* crispant (bottom). Microglia reduction observed in transient *il-34* crispants shown by neutral red staining (microglia, white arrows), phenocopying previously described stable *il-34* mutants[128]. **g** Quantification of macrophage infiltration indicates a significant reduction at 8-10 hpi in 4 dpf transient *il-34* deficient crispants. **h** Stable *il34* mutants derived from a heterozygous incross show either few or no macrophages in the liver after brain-LPS injection at 16-24 hpi similar to baseline brain-water injected animals, showing a much stronger effect in eliminating macrophage infiltration than in the partial gene knockout in transient *il-34* crispants. Statistical significance was determined by a two-tailed t-test coupled with a F-test validating equal variances for two-way comparisons, and Kruskal-Wallis multiple comparisons test for three-way comparisons in 3d and 3h (shown by the top bar) followed by corrected two-way tests if the multiple comparisons test was significant. *, p <0.05; **, p <0.01; ns, not significant; data points in scatter plots represent independent samples or animals.

**Figure 4.**
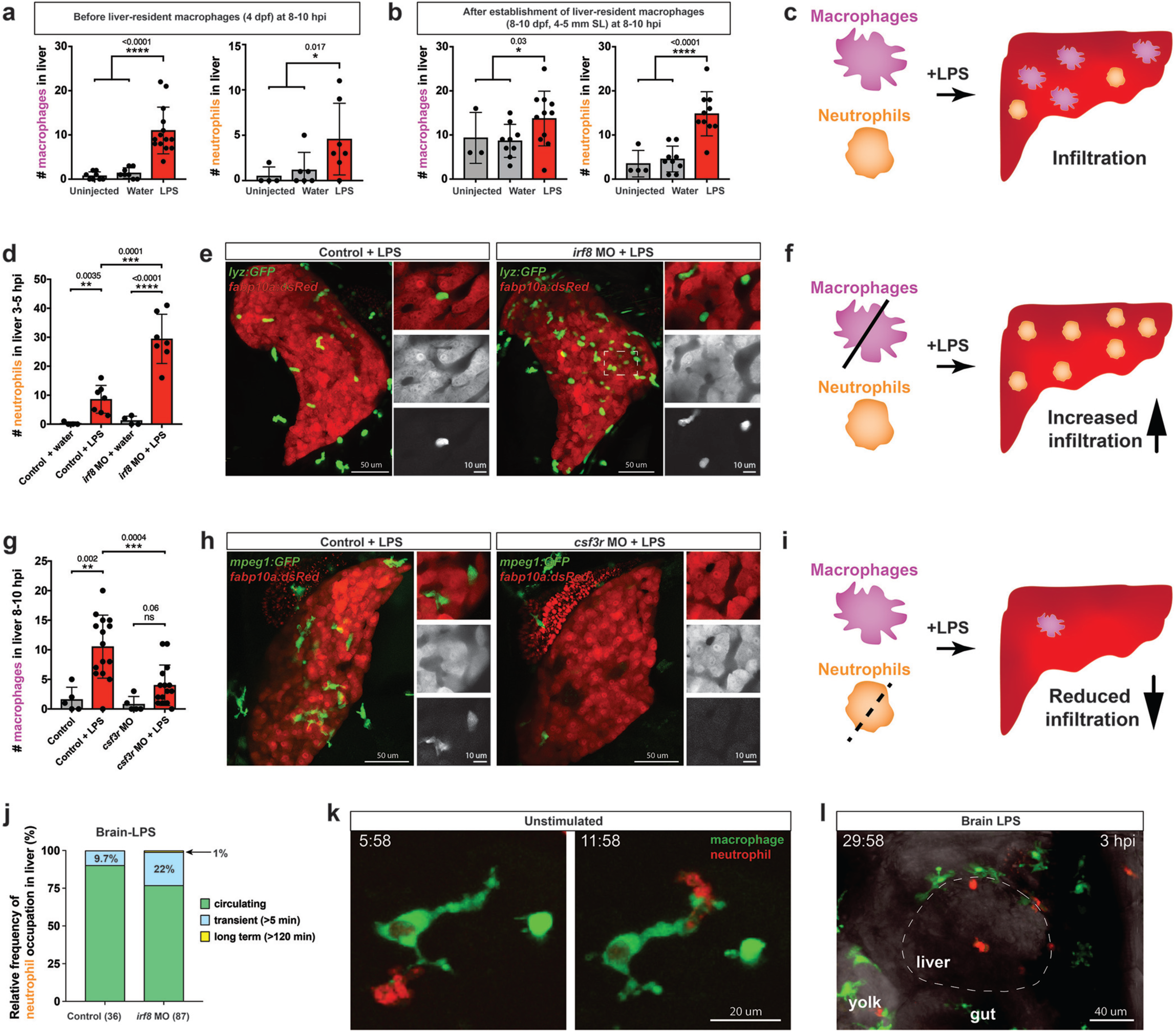
Neutrophils and macrophages coordinate to infiltrate the liver during a systemic inflammatory response. **a-b** Total number of macrophages and neutrophils in entire liver in control (uninjected and brain-water injected) and brain-LPS challenged animals before and after development of liver-resident macrophages (Kupffer cells) at 4 dpf and 8-10 dpf, respectively. **c** Diagram showing typical immune infiltration after LPS addition in wildtype animals. **d-e** Effects of macrophage ablation by *irf8* knockdown on neutrophil numbers in the liver 3.5-5 hours after brain LPS injection at 4 dpf. **d** Quantification of neutrophil numbers, **e** for control and irf8 morphants injected with LPS in brain at 4 dpf: left, 3D volume of whole liver. Visibly more neutrophil infiltration in *irf8* morphants than controls. Right, single z-plane images showing high magnification of a small region in liver shown on left. Individual infiltrated neutrophils were prevalent in gaps between hepatocytes, presumably sinusoids, after LPS injection. Top, merged channels; middle, hepatocytes (DsRed+); and bottom, neutrophils (GFP+). **f** Diagram illustrating an increase in neutrophil infiltration after macrophage ablation. **g-h** Depletion of neutrophils using the *csf3r* morpholino reduces macrophage infiltration compared with control LPS injections 8-10 hours after brain LPS injection at 4 dpf. **g** Quantification of macrophage numbers. **i** For control and *csf3r* morphants: left, 3D volume of whole liver; right, single z-plane showing a high magnification of a small region in liver shown on left. Significantly fewer macrophages were observed in the liver after neutrophil ablation. **i** Diagram summarizing the effect of neutrophil reduction. **j** Relative frequency of each type of neutrophil occupation in the liver after brain-LPS injection, as determined by *in vivo* time-lapse imaging. **k** Representative 3D images of normal macrophage and neutrophil interactions around the liver at 4 dpf (corresponding to Supplementary Movie 9). **l** 3D image of macrophage and neutrophil interactions after brain-LPS injection at 3 hpi at 4 dpf showing entry of neutrophils into liver prior to macrophages (corresponding to Supplementary Movie 8). Statistical significance was determined by a two-tailed t-test and with Welch’s correction for unequal variances as determined by a F-test. MO, morpholino. Each data point in scatter plots represents an independent animal. Transgenes used: *mpeg1:GFP* for macrophages, *lyz:GFP* for neutrophils, and *fabp10a:DsRed* for hepatocytes.

**Figure 5.**
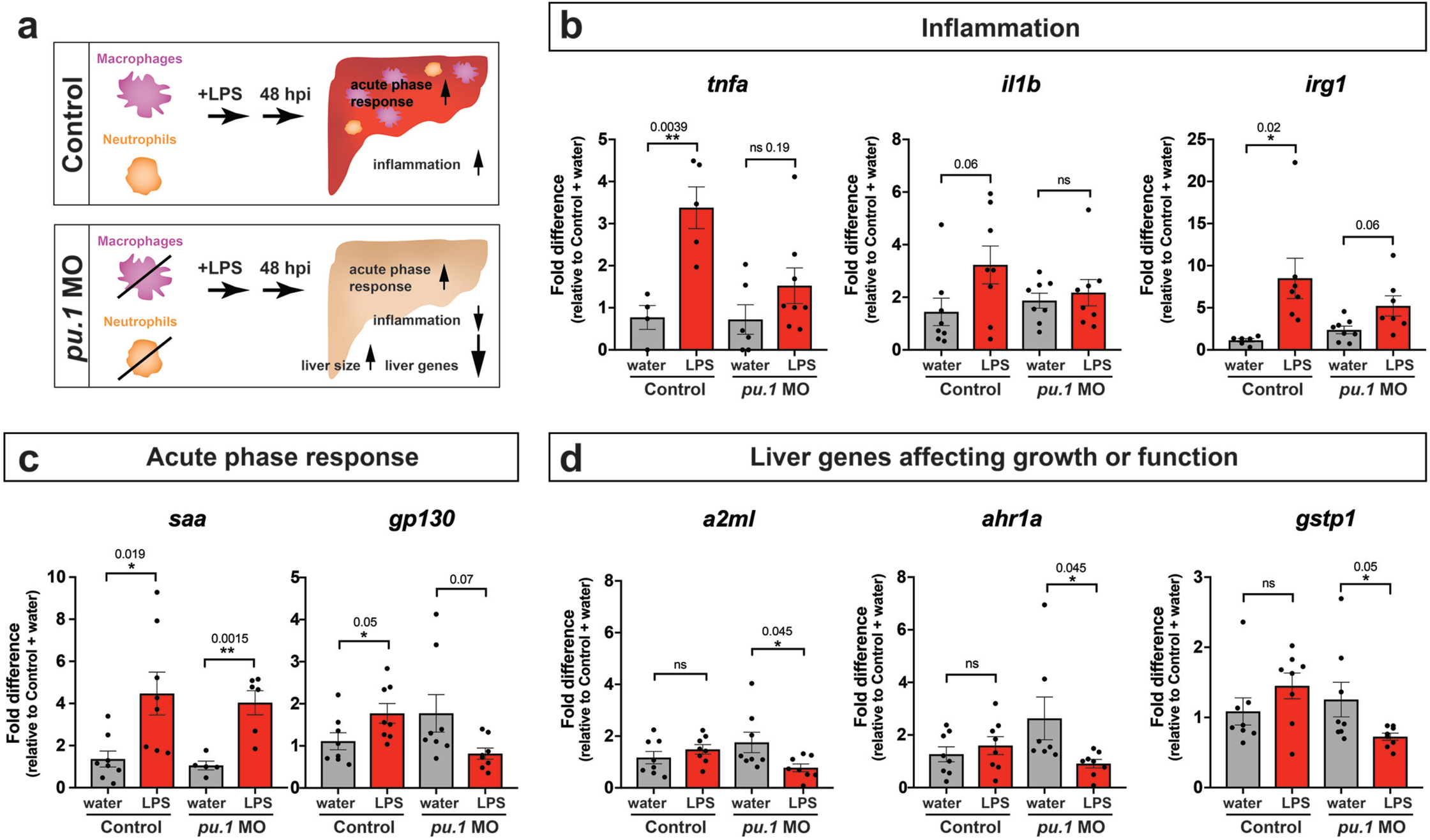
Eliminating myeloid cells disrupts hepatic response to systemic inflammation and causes transcriptional programmatic changes. **a** Schematic illustrates the impact of myeloid ablation by *pu*.*1* morpholino on hepatic response to brain LPS after 48 hours compared with the typical normal controls. **b-d** qPCR analysis was conducted on individual larvae 48 hours after either water vehicle or LPS injected in the brain in pu.1 morphants or control animals. **b** Relative expression of inflammation genes (*tnfa, il1b, irg1*) indicates decreased or reversal of inflammation in myeloid deficient animals while the inflammatory gene expressions remain highly elevated in wildtype animals after brain-LPS activation. **c** Acute phase response (APR) appears intact even in myeloid-deficient *pu*.*1* morphants after brain-LPS perturbation as saa, one of its major markers largely expressed by the liver, remains highly upregulated, although not *gp130* but it is not strictly an APR gene. **d** Three genes expressed in the liver that affect liver growth and function in zebrafish (*a2ml, ahr1a, gstp1*) were found to be all significantly downregulated in myeloid-deficient animals but not control wildtype at 48 hpi. Scatter plots show individual animals. Statistical significance was determined by a two-tailed t-test and with Welch’s correction for datasets with unequal variances. ns, not significant.

**Figure 6.**
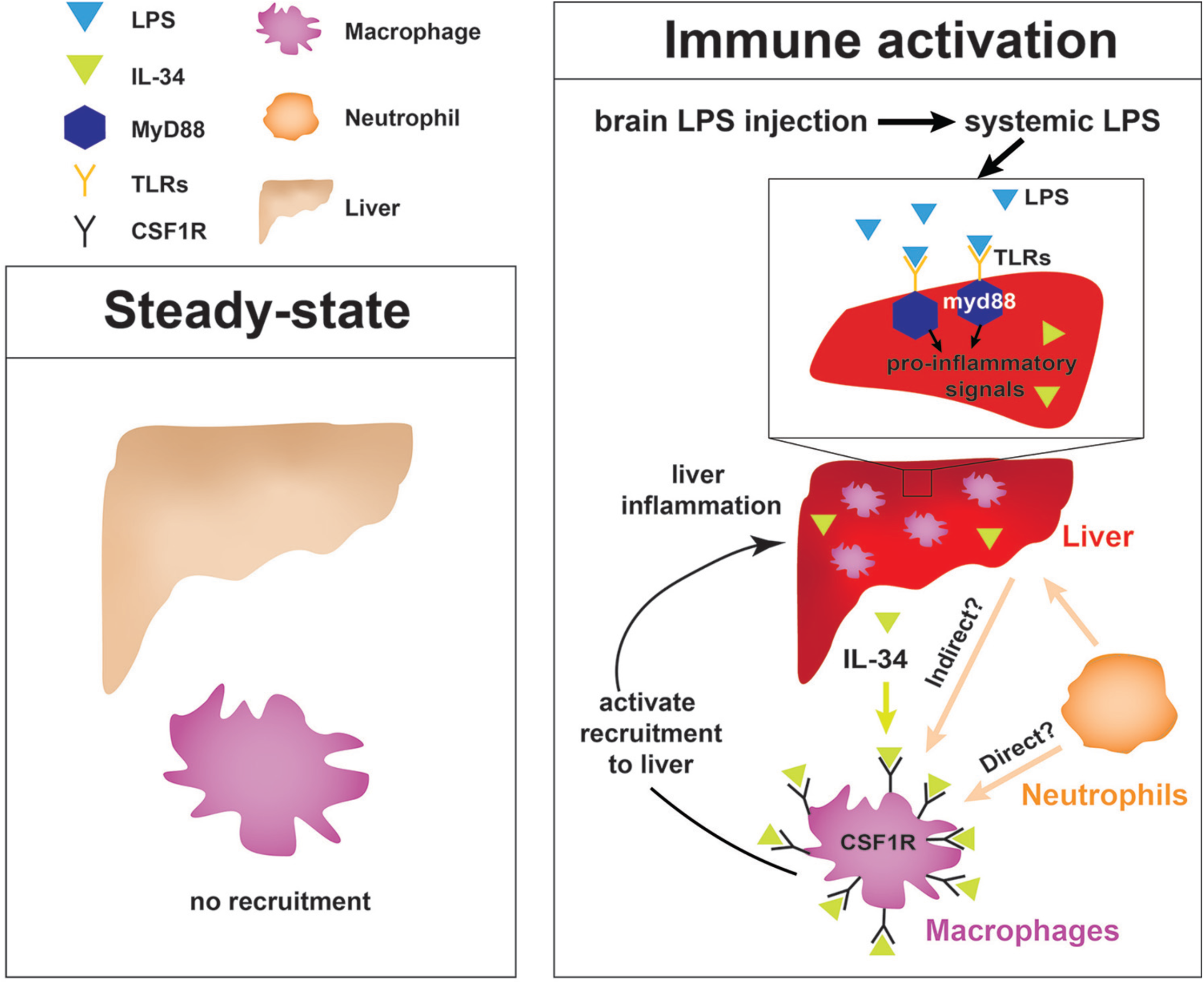
Current model for hepatic response to brain LPS drainage to the periphery. Diagram represents events happening prior to Kupffer cell development in the 4 dpf zebrafish. Left, at steady-state, no peripheral macrophages normally migrate into the liver. Right, recruitment of macrophages and monocytes into the liver after immune activation. Brain injection leads to systemic distribution of LPS that prominently induces a hepatic response whereby macrophages/monocytes are recruited, and genes conferring inflammation and acute phase response are highly upregulated. Recruitment of macrophages/monocytes into the liver requires MyD88, a common adaptor protein for TLRs that recognize LPS. In addition, IL-34 presumably secreted by the liver and downstream of MyD88 signaling may act as a chemoattractant to macrophages/monocytes expressing CSF1R. Macrophage recruitment also depends on yet unknown direct or indirect signaling from neutrophils, which infiltrate the liver first subsequent to drainage of LPS into circulation after brain LPS injection.

To characterize LPS at the tissue and cellular level along its outflow path, we used high power confocal imaging on the same LPS-injected zebrafish which were tracked by stereomicroscopy (Fig. 2 and Supplementary Movie 7) to carry out further imaging at later timepoints. Interestingly, LPS molecules were taken up by brain lymphatic endothelial cells (blec), also known as fluorescent granular perithelial cells (FGPs), as well as by facial and trunk lymphatic vessels as shown by co-localization of LPS with the lymphatic reporter *mrc1a:GFP* at 1.5 hpi [36](Supplementary Figs. 6 and 7), suggesting LPS exited through these lymphatic structures. These results were consistent with the previous LPS tracing within the first hour after injection (Fig. 2 and Supplementary Movie 7), but shown more definitively by high-resolution confocal imaging. By contrast, LPS were not localized within the CNS vasculature using the endothelial reporter *kdrl:mCherry*, but only in the peripheral blood vessels (Supplementary Fig. 7), indicating that transport of LPS did not likely result from a disruption of the BBB or a direct transit through brain blood vessels. Tracing the movement of LPS from the brain to the periphery also revealed its transient flow through the liver sinusoids prior to immune cell infiltration (Fig. 2, Supplementary Movie 5). However, LPS did not appear to accumulate or bind to cellular structures within the liver as we did not detect LPS there (Fig. 2b, Supplementary Fig. 7 and Supplementary Movie 5), suggesting either the transient exposure of hepatic cells to LPS, or yet unknown extrahepatic signals trigger the hepatic response to systemic LPS. Taken together, LPS appeared to enter circulation via a mechanism directed by the lymphatics for clearing away excess substances from brain parenchymal and surrounding interstitial fluids (Fig. 2e).

In light of the broad LPS distribution, we sought to functionally test whether the liver response after brain-LPS activation was due to systemic LPS. To create a systemic LPS condition akin to mammalian models of sepsis or endotoxemia[37, 38], we directly injected LPS intravenously at the caudal plexus into the bloodstream and compared its resulting peripheral response to that after brain-LPS injection (Fig. 2). These injections resulted in different outcomes for microglial activation at 6 hpi, whereby brain-LPS led to a strong activation of microglia but intravenous (IV)- LPS did not (Fig. 2f). Both routes, however, led to the same robust macrophage infiltration of the liver (Fig. 2g). To determine if circulation was required for the liver response after brain-LPS injection, we used a morpholino to knockdown *tnnt2*, a cardiac muscle troponin T gene, a well-established reagent for blocking circulation[39] (Supplementary Fig. 9). We indeed found that inhibiting circulation prevented macrophage infiltration into liver after brain-LPS injection (Supplementary Fig. 9). Taken together, several lines of evidence show that circulating LPS that causes systemic inflammation was driving macrophage infiltration into the liver after brain-LPS injection: 1) systemic distribution of LPS, 2) sufficiency of IV-LPS injection to cause liver infiltration, and 3) blocking circulation prevented macrophage infiltration of the liver.

### Macrophage recruitment to the liver is a MyD88-dependent inflammatory response requiring IL-34 pathway for migration

We next tested the effects of anti-inflammatory drugs on liver response after brain-LPS microinjection to address whether systemic inflammation was indeed responsible for the macrophage recruitment into the liver. We screened five small-molecule drugs (GW2580, 17-DMAG, celastrol, Bay 11-7082, and dexamethasone) known to effectively curb inflammation by attenuating NF-kB mediated transcription or activating glucocorticoid functions in zebrafish and other systems[40-49]. These small molecules are known to act through different mechanisms: GW2580, a selective inhibitor of cFMS kinase that blocks the receptor tyrosine kinase CSF1R function which can prevent NF-kB activation [50]; 17-DMAG (a water-soluble geldanamycin analog) and celastrol, both potent inhibitors of the heat-shock protein Hsp90 that cause disruption or degradation of its target proteins, including the NF-kB protein complex[41, 51]; Bay 11-7082, an inhibitor of E2 ubiquitin (Ub) conjugating enzymes, which target NF-kB inhibitor, IkB-alpha, for proteasomal degradation[49]; and dexamethasone, an agonist of the glucocorticoid receptor (GR) that activates a negative feedback mechanism to reduce inflammation[47]. We first assessed the effects these small molecules had on macrophage infiltration into the liver after brain-LPS microinjection using whole-mount RNA *in situ* hybridization with the macrophage marker *mfap4* (Supplementary Fig. 10). As positive controls for liver infiltration, we used brain-LPS injected animals that were untreated (water only) for comparison with water-reconstituted drugs (17-DMAG and dexamethasone), and treated with only DMSO for comparison with DMSO-reconstituted drugs (GW2580, celastrol, and Bay 11-7082). Untreated animals without brain microinjection were also used as negative controls. By *in situ* analysis, we found that Bay 11-7082 and dexamethasone substantially reduced the frequency of macrophage infiltration after brain-LPS injection compared with control groups (Supplementary Fig. 10 a, b). These effects were further validated by *in vivo* imaging of macrophage recruitment into the liver, which enabled a precise macrophage count at a high cellular resolution (Supplementary Fig. 10 c,d). These results indicated that macrophage infiltration into the liver after brain-LPS stimulation can be prevented by suppressing inflammation via mechanisms inhibiting the NF-kB pathway or activating the glucocorticoid signaling.

To examine possible components affiliated with the NF-kB pathway that may drive macrophage recruitment into the liver, we examined whether an intracellular signal adaptor protein myeloid differentiation protein-88 (MyD88) of Toll-like receptors (TLRs) known for recognizing LPS and mediating cytokine production was essential[52]. We employed an effective *myd88* specific splice-blocking morpholino to knockdown *myd88* as previously described[53] during the liver response to brain-LPS stimulation (Fig. 3 a, b). The efficacy of *myd88* morpholino to mediate splice-blocking was confirmed by RT-PCR analysis (Fig. 3c). By *in vivo* imaging, we found that the number of liver-infiltrating macrophages after brain-LPS microinjection was significantly reduced when *myd88* function was disrupted (Fig. 3 a, b). To fully validate these results, we performed the brain-LPS and control injections in genetic *myd88* null mutants and their siblings derived from a heterozygous incross (Fig. 3d). *myd88* mutants showed either few or no macrophages in the liver after brain-LPS injection at 16-24 hpi similar to baseline brain-water injected animals (Fig. 3d), demonstrating a much stronger effect in reversing macrophage infiltration than the partial *myd88* knockdown by morpholinos. These results show that the inflammatory liver response depended on the MyD88 pathway and are consistent with LPS-triggered inflammation as the driver of macrophage infiltration into the liver.

As possible hepatic signals that recruit macrophages into the liver during systemic inflammation, we examined whether the interleukin-34 (IL-34) and colony stimulating factor-1 (CSF-1) that share a common receptor CSF1R had a role. IL-34 and CSF-1 are known to mediate various functions of macrophages including inflammatory processes and promoting production of pro-inflammatory chemokines[50, 54, 55]. They have recently been shown to be required for macrophage migration and colonization of the brain to form microglia in zebrafish[56-58]. Interestingly, engineering an artificial expression of *il-34* in hepatocytes has been shown to be able to recruit macrophages to the liver in zebrafish, but the physiological relevance was not known[59]. In light of these previous studies, *il-34* and the two zebrafish orthologs of CSF-1 gene (*csf1a* and *csf1b*) were strong candidates for attracting macrophages to the liver after brain-LPS injection. To examine this possibility, we first determined whether these genes (*il-34, csf1a*, and *csf1b*) were upregulated in the liver after brain-LPS injection using quantitative PCR (qPCR) analysis on liver-specific and body-minus-liver tissues compared with control animals without the brain microinjection (Fig. 3e). We found that while *csf1a* and *csf1b* were either not detected or had no difference in the liver with or without LPS injection, *il-34* was significantly upregulated in the liver after brain-LPS microinjection (Fig. 3e). This upregulation was specific to the liver as *il-34* was not elevated in the body-minus-liver tissue after brain-LPS injection (Fig. 3e). To test if *il-34* was a required cytokine for recruiting macrophages in our experimental paradigm, we used CRISPR/Cas9 targeted mutagenesis as previously described[28, 56] to disrupt *il-34* function to assess the immune infiltration (Fig. 3 f-g, Supplementary Fig. 11). These *il-34* targeting gRNA/Cas9 injected animals are coined as “crispants”. To verify the efficacy of the gene knockdown in transient *il-34* crispants, we determined whether they phenocopied a reduced microglia phenotype recently described in *il-34* stable mutants[57]. Indeed we found about 40% of the transient *il-34* crispants to have highly decreased microglial numbers (Fig. 3f). By Sanger sequencing analysis, we verified that these crispants induced a high frequency of frameshift indels altering reading frames and introducing early stop codons in the *il-34* locus (Supplementary Fig. 11). We found that transient *il-34* crispants after brain-LPS injection indeed had a significantly reduced number of infiltrating macrophages compared with controls (no crispant) after brain-LPS injections (Fig. 3g). To confirm these results, we also tested the effects of brain injections in stable *il-34* null mutants and their control siblings derived from a heterozygous incross (Fig. 3h). *il-34* mutants had either few or no macrophages in the liver after brain-LPS injection at 16-24 hpi similar to baseline brain-water injected animals (Fig. 3h), providing clear evidence that *il-34* is essential in recruiting macrophages to the liver after brain-LPS activation. Taken together, these results showed that macrophage infiltration into the liver was driven by inflammatory processes that require the *myd88* pathway as well as *il-34* signaling likely coming from the liver.

### Liver infiltration by macrophages is driven and coordinated by neutrophils

Since inflammation is typically a concerted response of the innate immune system, the liver response after brain-LPS stimulation may involve other immune cells besides macrophages. To examine this possibility, we investigated whether neutrophils, the other functional leukocytes prominent at early larval zebrafish stages[60, 61], could participate in infiltrating the liver along with macrophages. Using live imaging in transgenic zebrafish at 4 dpf prior to Kupffer cell development, we quantified the numbers of neutrophils and macrophages in the liver 8-10 hpi with control water or LPS, as well as in uninjected controls (Fig. 4a). We performed the same brain injection experiments also at 8-10 dpf after Kupffer cell establishment (Fig. 4b). Our data interestingly showed that neutrophils also significantly infiltrated the liver after brain-LPS but not in the control brain-water and uninjected groups. Similar to macrophages, neutrophils infiltrated the liver irrespective of the presence of Kupffer cells (Fig. 4 a-c). To test whether liver infiltration would still occur at juvenile adult stages, when the liver, blood and lymphatic vasculature, and brain structures are fully mature, we conducted brain tectum injections in 1 month-old zebrafish which had a standard length of 0.8-1 cm (Supplementary Fig. 12). We found at 16-18 hpi that the numbers of macrophages and neutrophils in the liver were significantly increased after LPS injection compared with water vehicle injections and uninjected controls (Supplementary Fig. 12), indicating that the liver infiltration persists even at young adult stages. These results implicate that the hepatic response to brain-LPS injection is independent of Kupffer cells, age, or maturity of the brain vasculature and architecture, and drainage of LPS from brain to periphery may still be evident in adulthood. They also raise the possibility that macrophages coordinate with neutrophils for moving into the liver, and signals other than from Kupffer cells can recruit these leukocytes into the liver during systemic inflammation induced by circulating LPS.

To interrogate a possible coordination between macrophages and neutrophils, we turned to gene perturbations to examine the functional consequence of eliminating one immune cell type on the other. We utilized a previously established morpholino targeting *irf8*, an essential transcription factor for macrophage formation, whose effect phenocopies the macrophage-lacking *irf8* genetic mutants for ablating all macrophages at embryonic and early larval stages[62, 63]. Assessment of neutrophil infiltration was conducted at 3-5 hpi, which is earlier than the timepoint for macrophage analysis, because we found neutrophils to infiltrate the liver first before macrophages (Fig. 4l, Supplementary Movie 8). Strikingly, a highly significant 3-fold increase of liver-infiltrating neutrophils was found in macrophage-ablated *irf8* morphants after brain-LPS injection at 4 dpf compared with the control brain-LPS animals, while brain-water injected controls had no neutrophil infiltration of liver with or without macrophages (Fig. 4d). To verify these findings, we used a complementary approach for macrophage depletion by using clodronate-containing liposomes to induce apoptosis in macrophages as previously described[64] (Supplementary Fig. 13). Significantly increased neutrophil infiltration of liver after brain-LPS injection in clodronate-mediated macrophage depletion was also found compared with the control response to brain-LPS (Supplementary Fig. 13). Both methods of macrophage depletion corroborate to show striking increases in neutrophil recruitment to the liver after brain-LPS microinjection. These results suggest that neutrophils may compensate for macrophage loss, or that they are normally constrained by macrophages in part possibly by being phagocytosed as they become impaired or die, as has been described in an immune response[65]. However, these infiltrating neutrophils do not appear to fully make up for macrophage functions as they do not exhibit the same dynamic movements or lasting occupation in the liver during the hepatic response to brain-LPS (Fig. 4j). Most of the infiltrating neutrophils are circulating through the liver (>80%) regardless of macrophage presence in the liver during brain-LPS response, albeit a larger fraction is transient (22%) and long-term (1%) after macrophage ablation (Fig. 4j). Our data therefore favors the latter explanation that macrophages may limit ongoing recruitment of inflammatory neutrophils.

Conversely, we employed a splice-blocking morpholino targeting the granulocyte colony-stimulating factor receptor (CSF3R/GCSFR) for reducing neutrophil numbers in zebrafish[66]. We verified by *in situ* gene expression pattern of the neutrophil marker, *mpx*, that *csf3r* morphants indeed had neutrophil numbers reduced on average ∼30% with a maximum of a 65% decrease (Supplementary Fig. 14). Also, by in *vivo* imaging of larval zebrafish carrying both macrophage and neutrophil reporters, the *csf3r* morphants showed no significant change in macrophage number, but a substantial neutrophil reduction (Supplementary Fig. 14). These results indicate *csf3r* knockdown does not impact baseline macrophages, consistent with neutrophil-specific reduction and functional defects shown in zebrafish *csf3r* mutants[66, 67]. In these neutrophil-reduced *csf3r* morphants after brain-LPS injection, we found significantly reduced macrophage infiltration of liver by *in situ* and live imaging analyses (Fig. 4 and Supplementary Fig. 14), indicating the requirement of normal neutrophil presence for liver infiltration. In agreement, constant and dynamic intermingling of these two immune cell types were found around the liver normally (Fig. 4k and Supplementary Movie 9). These results taken together show that during liver inflammation, macrophages and neutrophils coordinate intimately, and macrophages are at least in part driven by neutrophils to enter the liver as they require neutrophils which infiltrate the liver first (Fig. 4l, Supplementary Movie 8).

### Immune infiltration of liver promotes inflammation and disrupts liver growth

The functional consequence of macrophage and neutrophil infiltration on the liver during a systemic inflammatory response remains unclear. To examine this, we assessed transcriptional changes comparing immune infiltration and lack thereof in the absence of macrophages and neutrophils. Previous studies have implicated the role of infiltrating innate immune cells in causing liver injury and inflammation[18] but whether this effect is conserved in this model of brain-triggered systemic inflammation is unknown. We examined by qPCR analysis a set of 8 genes after brain-LPS injection with or without innate immune cells compared with the control brain-water injected group to gauge the long-term effect of immune infiltration at 48 hpi. Disruption in liver homeostasis can be defined by an upregulation of genes associated with inflammation (such as pro-inflammatory cytokines *tnfa* and *il1b*). Liver under stress, trauma and inflammation is also characterized by activating an acute phase response[68] (such as serum amyloid A (*saa*) and interleukin 6 signal transducer (*il6st/gp130*)). Disrupted liver can also exhibit altered expression level of genes associated with liver growth and function (such as alpha-2-macroglobulin-like (*a2ml*)[69], aryl hydrocarbon receptor 1a (*ahr1a*)[70], and glutathione S-transferase pi 1 (*gstp1*) in zebrafish)[71]. To eliminate myeloid cells, we targeted an essential myeloid transcription factor *pu*.*1/spi1b* using an established translation-blocking morpholino previously shown to eliminate macrophages/microglia and neutrophils[62]. The efficacy of *pu*.*1* morpholino was validated by finding a complete loss of brain macrophages (microglia) in *pu*.*1* morphants using a neutral red staining assay (n=4/4) as previously described[72]. The qPCR results indicated that in the control situation with the full complement of macrophages and neutrophils at two days after brain-LPS injection in the 4 dpf zebrafish larvae, all genes associated with inflammation and acute phase response were significantly upregulated, while genes known to affect zebrafish liver development and function were not changed (Fig. 5). By contrast, depletion of myeloid cells in *pu*.*1* morphants resulted in no significant upregulation in all inflammation genes analyzed (Fig. 5b), indicating that the innate immune cells were responsible for the upregulation of inflammatory genes.

Furthermore, *saa* remained highly upregulated as in control brain-LPS animals, but liver genes associated with growth or function were significantly downregulated compared with water vehicle injected morphants (Fig. 5). Since *saa* transcript level was not modified by an absence of macrophages, we further asked whether *saa* may be dispensable for macrophage infiltration. Comparing brain-LPS injections in *saa*^-/-^ mutants side-by-side with injected control siblings showed no difference between the genotypes (Supplementary Fig. 15), indicating that *saa* is not required for macrophage infiltration into the liver upon LPS activation. The downregulation of liver genes in myeloid cell-depleted animals after brain-LPS injection raised the question as to whether this reflected an actual change in liver growth. To address this, we used *in vivo* confocal 3D imaging to capture the whole liver at 48 hpi to measure liver size by volume in the different backgrounds and treatments (Supplementary Fig. 16). We found a decrease in liver size after LPS injection compared with water vehicle injection in both baseline and control-MO groups, but, interestingly, increased liver sizes in LPS-injected myeloid-lacking *pu*.*1* morphants (Supplementary Fig. 16), indicating liver growth was likely impeded by immune cell infiltration during inflammation. Since the expression of liver genes decreased but the liver size increased, these liver genes may not be associated with growth but rather function. These results indicate that the liver infiltration by macrophages and neutrophils promoted inflammation and disrupted liver growth, but had no effect on the acute phase response gene *saa*, which was not essential for the infiltration (Supplementary Fig. 15). Taken together, they raise the possibility that immune cell infiltration leads to liver resources and functions being redirected from normal developmental growth to a full-fledged inflammatory response.

## Discussion

### Drainage of macromolecules from brain to body engages the periphery to respond to central changes

As a rapid means by which the brain can engage the periphery in response to possible danger, our study reveals how inflammatory molecules introduced into the brain parenchyma can circulate outside of the brain to initiate a robust hepatic response in zebrafish (Fig. 6). It remains to be explored whether the transport of molecules or substances outside of the brain even at trace levels resulting from CNS trauma or injury can explain at least to some degree the associated hepatic damage described in mammals[24, 73, 74]. The most likely route through which macromolecules can drain out of the brain is through the lymphatic system that removes interstitial fluids, waste products, and immune cells in zebrafish and mammals[74-76]. Recent new knowledge in meningeal lymphatics and glymphatics indicate that macromolecules in the brain can flow from the parenchymal interstitial space into the cerebrospinal fluid (CSF) which drains into the lymphatic vessels or directly into the major veins (sinuses) to enter general circulation[74, 75, 77, 78]. In support of this, previous studies using a number of mammalian species (cat, rabbit, dog, and rat) have shown that injection of tracers at 5-100 ul volume such as horseradish peroxidase, India ink or dextran blue into the CSF or brain parenchyma gets drained rapidly into cervical lymph nodes within seconds as well as in entire vasculature within minutes[79-84], while radiolabeled albumin microinjected into the brain parenchyma of different brain regions appear rapidly in the CSF and within 4 hours can be found in cervical lymph nodes and the common carotid arteries[84, 85]. Furthermore, metabolites of glucose from the brain are found to be released in the cervical lymph nodes[86], supporting the possibility that drainage of endogenous macromolecules from brain to blood circulation can also happen. Taken together, studies in mammals indicate the ability for molecules to transport from the parenchyma or CSF-filled ventricles into general circulation either through the lymphatics, or blood vessels directly. However, the physiological impact of brain drainage on the periphery had not been investigated, nor directly visualized. Our study demonstrates that such drainage can be directly tracked *in vivo* to induce a robust hepatic response defined by an active recruitment of inflammatory macrophages and neutrophils into the liver.

Similarly, we found that brain tectum microinjection in zebrafish leads to effects comparable to that known of mammalian intracerebroventricular, intracisternal, and intraparenchymal injections [35, 77, 78], such that the injected LPS in the parenchyma flows into the CSF compartment (or ventricles) rapidly and likely via bulk flow (Fig. 2 and Supplementary Movie 7), followed by clearance of LPS and ISF predominantly by lymphatic drainage (Fig. 2a-b, Supplementary Figs. 6 and 7). After brain-LPS injection, we found a low-level spread of LPS/CSF from the hindbrain ventricle into the brain parenchyma and surrounding interstitial space prior to LPS accumulation in brain lymphatic cells and passage through the extracranial lymphatic vessels. Whether CSF flows to the brain more widely than the major perivascular spaces upon intracranial injection remains controversial in mice [77, 78], although this appears to occur easily in the larval zebrafish perhaps due to its small brain size conducive to simple diffusion or bulk flow of fluids. Previous work in mice have shown that injected tracers in the subarachnoid CSF or lateral ventricle can enter the brain parenchyma quickly in less than 30 minutes, and get cleared through paravascular or perineural pathways via the lymphatic system similarly to intraparenchymal injections [77, 78], indicating the process of drainage from the brain parenchyma may be conserved between zebrafish and mammals. On average, at about 13 minutes after tectum microinjection of fluorescent LPS in larval zebrafish, we detected initial signals of the LPS in circulation, albeit at a baseline intensity, far less than that at the injection site and in the ventricles. This is about two times faster than the transit time for a lateral ventricle injection in mice with a fluorescent tracer to reach blood circulation (around 25 minutes)[78], which is still relatively fast. Two key traits that may endow larval zebrafish a faster rate of drainage of solutes from brain ventricle or parenchyma than in rodents may be: 1) the short physical distances between anywhere in the parenchyma and the nearest ventricle and vasculature, which would span no farther than the width of the brain, approximately a few hundred microns, enabling simple diffusion as a primary mode of transport [10], and 2) the lack of lymph nodes, based on the prevailing understanding of the zebrafish anatomy [87, 88], which are lymphatic structures throughout the lymphatic network in mammals that capture and filter the fluid that exits the CNS before joining circulation. The content and concentration of the molecules leaving the CNS that reach blood circulation may be limited by lymph nodes in mammals, but these restrictions may be more lenient in zebrafish due to a lack of these structures. Instead of lymph nodes, we found that the intraparenchymally-injected LPS in larval zebrafish after spreading in the parenchymal and surrounding interstitial space is engulfed by brain lymphatic endothelial cells (blec), which are non-lumenized [89-91], as well as collected by the facial and trunk lymphatic vessels (Supplementary Figs. 6 and 7). These non-lumenized blec also exist in mouse [91], but their contribution to the clearance of solutes from the CSF/ISF is yet unclear. Interestingly, while facial and trunk lymphatic vessels surrounding the brain were filled with LPS by 1.5 hpi, localization of LPS within the brain blood vessels was not observed (Supplementary Figs. 6 and 7), indicating the main route by which LPS exit the CNS to enter general circulation was via the lymphatic tracts. Additionally, in both zebrafish and mammals, it is also possible that macromolecules to some degree get eliminated by ingestion by perivascular and other phagocytic cells before they drain out to the periphery through the lymphatic-vascular system.

Whether the process of drainage differs during developmental stages when the tissue boundaries and structures of the brain are less mature, we provide evidence that the peripheral hepatic response to brain-LPS injection may be age-independent in zebrafish. The hepatic response was observed from early to late larval stages, and also in juvenile adults, suggesting passage of inflammatory molecules occurs even after establishment of a fully mature brain architecture (including the blood-brain and blood-CSF barriers). While this study focused on the early larval stage at 4 dpf as it offers a timepoint at which infiltrating cells can be clearly assessed without the presence of resident macrophages (Kupffer cells), zebrafish at this age are functionally mature with established functional blood-brain barriers and ventricles with choroid plexus and CSF[25-27]. Regardless of the route or quantities at which macromolecules can drain out from the brain, our results implicate the process of drainage from brain as an effective mechanism by which the brain can quickly transmit molecular information to the periphery to engage a prompt hepatic response to potential CNS threats.

### Responsiveness of liver to systemic inflammation irrespective of Kupffer cells

Our data demonstrate that the most striking effect of LPS injection into the brain parenchyma was the recruitment of macrophages into the liver in the zebrafish model (Fig. 6). In agreement with the liver being most susceptible to systemic inflammation, previous studies using intravenous injection of the endotoxin LPS in different mammalian models have shown an immediate localization of LPS to the liver and less to other organs by tracking radiolabeled LPS[92-95]. Prior to our work, it was not known that introduction of endotoxin into the brain parenchyma can result in the same impact on the liver as that from a systemic administration. These findings beg the question as to how the liver is the predominant and conserved target of circulating LPS and systemic inflammation in vertebrates from zebrafish to mammals. One overarching explanation could simply be related to the sheer blood volume coming from the hepatic artery and portal vein [11] that the liver constantly processes, making it particularly sensitive to inflammatory cues in circulation.

Kupffer cells have been considered to be key immune cells responsible for producing cytokines and chemokines to recruit neutrophils and other leukocytes that may further propagate systemic inflammation[14, 96]. In fact, tracking of radiolabeled LPS has shown that circulating LPS was mostly taken up by Kupffer cells[92, 93], implicating these cells as the main sink for endotoxins or microbes. By contrast, our study shows that the liver is capable of responding to systemic inflammation and recruiting neutrophils first and then macrophages, independent of Kupffer cells. We found that starting in the larval zebrafish at 4 dpf when the liver is already functional and differentiated with diverse hepatic cell types including hepatocytes, sinusoidal endothelial cells, biliary cells, and stellate cells[97], but before Kupffer cells are established, other hepatic cells besides the Kupffer cells are capable of signaling to recruit peripheral leukocytes into the liver. Furthermore, while the gut may provide a source of stimulating molecules through the portal vein, we show that the brain is another source of inflammatory cues that can result in immune infiltration of the liver that was not previously appreciated.

### IL-34 pathway in mediating immune infiltration of liver triggered by systemic inflammation

The receptor tyrosine kinase CSF1R (also known as Fms) pathway regulates migration, differentiation, proliferation, and survival at varying degrees of different tissue-resident macrophages and monocytes in mammals and zebrafish[56, 57]. Recent studies have implicated a role for this pathway in macrophage recruitment in disease, including the assembly of tumor-associated macrophages in various cancers[98-102]. Of particular interest IL-34 is elevated and implicated in the inflammation process of several inflammatory diseases including rheumatoid arthritis, inflammatory bowel disease, and Sjogren’s syndrome[103-107]. Interestingly, among the three known ligands of the zebrafish homolog of the receptor tyrosine kinase CSF1R, namely *csf1a, csf1b*, and *il-34*, we found that *il-34* was singly upregulated in the liver upon brain-LPS microinjection. Moreover, we showed that disruption of *il-34*, either by antisense splice-blocking morpholinos or a stable loss-of-function genetic mutation, significantly eliminated macrophage infiltration of liver induced by brain-LPS microinjection (Fig. 3). The requirement for *il-34* signaling presumably from the liver may be coupled with *myd88* known to act downstream of TLR signaling upon LPS recognition, as *myd88* is also highly expressed in the normal larval zebrafish liver [108]. Whether these pathways act only in the liver, other cell types, or both, and whether other signaling mechanisms are involved, including CXCR3-CXCL11 known to affect macrophage chemotaxis to infection sites in human and zebrafish [109], remain to be explored. Our data indicate an essential role for *il-34* in recruitment of macrophages into the liver, implicating a new function for the *il-34/csf1r* pathway in conjunction with *myd88* dependent mechanisms during hepatic inflammation.

### Coordination of the innate immune system during hepatic response to systemic inflammation

Chronic active infiltration of leukocytes into the liver can cause liver damage and subsequent progression to fibrosis, cirrhosis or liver cancer [18-20]. This is a common feature shared among liver diseases caused by infection, toxic insults, or autoimmunity[17]. Using zebrafish, we found that macrophages and neutrophils were co-dependent in the process of infiltrating the liver after brain microinjection of LPS. We further showed by *in vivo* time-lapse imaging an early recruitment of neutrophils in the first few hours was followed by migration or circulation of peripheral macrophages into the liver. This is consistent with previous studies showing early neutrophil recruitment followed by monocyte-derived macrophages in other contexts of inflammation[110-112]. While secreted granule proteins from neutrophils that have already infiltrated the liver may directly recruit macrophages[113, 114] in our experimental platform of liver infiltration, we cannot exclude the possibility that indirect effects by which neutrophils alter vascular permeability or signaling from endothelial or other hepatic cell types actually direct macrophage recruitment into the liver.

Conversely, the impact of monocytes and macrophages on neutrophil activity remains less understood. Using two complementary approaches to deplete macrophages either by *irf8* genetic deficiency or liposomal clodronate treatment in zebrafish, we found that macrophages may be important to limit neutrophil infiltration into the liver upon brain drainage of LPS. An excessive level of neutrophil infiltration may lead to a heightened release of toxic metabolic products and proteolytic enzymes[65], thereby causing more damage to the liver. Macrophages in the liver are known to remove apoptotic or impaired neutrophils by phagocytosis, which in turn can modulate their own signaling based on the receptor(s) used during neutrophil clearance[113]. Since deficiency in *irf8* causes an elimination of macrophages but also an increase in baseline neutrophil number [63], we cannot completely rule out that a larger neutrophil pool may contribute to the significant increase in neutrophil infiltration after brain-LPS injection in *irf8* morphants. Nonetheless, the agreement in effect from two distinct methods of macrophage ablation, and the observed intimate macrophage-neutrophil contacts in the hepatic region strongly support reciprocal interactions between macrophages and neutrophils.

### Perspective on the brain-liver connection

Our study shows that the drainage following microinjection of LPS into the brain robustly caused peripheral macrophages and monocytes to infiltrate the liver, a process mediated by an upregulation of the cytokine IL-34 in the liver presumably downstream of activating the Toll-like receptor adaptor protein MyD88, and signaling from neutrophils. Treatment with a subset of anti-inflammatory drugs indicates that glucocorticoid activity or inhibiting NF-kB activation suppresses immune cell infiltration during hepatic response to systemic inflammation. These results prompt intriguing questions on how a hepatic response to a central disruption may reflect the presence and severity of CNS inflammation; whether a hepatic response contributes to the recovery or disruption of brain homeostasis; and to what extent does drainage of harmful or inflammatory molecules from brain to circulation explains for the peripheral problems associated with primarily CNS pathologies. Understanding mechanisms regulating immune infiltration of the liver and brain-periphery interactions can offer new approaches for modulating central inflammation while limiting damage elsewhere in the body.

## Materials and methods

### Zebrafish

Embryos from wild-type (TL and AB), mutant and transgenic backgrounds: *il34*^*re03*^[115], *myd88*^*b1354*^[116], *saa*^*rdu60*^[117], *mpeg1:EGFP [118], lyz:GFP[119], lyz:mCherry; cmlc2:GFP*[120], *fabp10a:DsRed[121], kdrl:mCherry-CAAX[122], mrc1a:egfp*^*y251*^*[36], mfap4:tdTomato[123], and nbt:DsRed[124]* were raised at 28.5°C and staged as described[125]. Stable transgenic *mpeg1:BFP* and *fabp10a:BFP* fish lines were generated using Tol2-mediated transgenesis based on cloned constructs combining the BFP coding sequencing (gift from Martin Distel) downstream of their respective regulatory sequences as published for *mpeg1(1*.*86 kb)[126]* and *fabp10a(2*.*8kb)[117]*. This study was carried out in accordance with the approval of UNC-Chapel Hill Institutional Animal Care and Use Committee (protocols 16-160 and 19-132).

### LPS and bacteria microinjections

Zebrafish larvae at 4-10 dpf and 1 month old were mounted dorsal side up for brain injections or on their sides for intravenous injections in 1-3% low-melting agarose. A pneumatic microinjector (WPI) with a fine capillary glass pipette was used to inject 1 nL for 4-10 dpf or 2 nL for 1 month old of stimulus into the targeted site (either tectum for brain injections, or caudal vein plexus for intravenous injections). Lipopolysaccharides (LPS) derived from 0111:B4 *E. coli* (L3024 Sigma) at 5 ng/nL, ultra-pure water, or live *Escherichia coli* cells were supplemented with fluorescently labeled dextran (Invitrogen, 10,000 MW at 1:100 dilution of a 5 ng/nL stock) for visualization. *E. coli* cells were prepared for injections as previously described[28]. Fluorescently tagged Alexa 594 conjugated LPS from 055:B5 *E. coli* (L23353 Sigma) at 5 ng/nL was used for brain microinjection at 1 nL per fish. All fish after microinjection are carefully monitored for normal health and behavior, and nearly all injected fish remain healthy and viable, showing no signs of overt change. These healthy post-injection fish are used for further experimentation.

### *In vivo* time-lapse and static confocal imaging

All time-lapse and static z-stack imaging were performed using a Nikon A1R+ hybrid galvano and resonant scanning confocal system equipped with an ultra-high speed A1-SHR scan head and controller. Images were obtained using an apochromat lambda 40x water immersion objective (NA 1.15) or a plan apochromat lambda 20x objective (NA 0.75). Z-steps at 1-2 um were taken at 40x and 3-5 um at 20x. Different stages of zebrafish were mounted on glass-bottom dishes using 1.5% low-melting agarose and submerged in fish water supplemented with 0.003% PTU to inhibit pigmentation. Dissected juvenile and adult liver tissues were mounted in fluoromount-G (Southern Biotech) for imaging.

### Liver cell counts

At larval stages 4-10 dpf, the whole liver in the live transgenic larvae was captured in a z-stack used for counting total number of macrophages and neutrophils in the liver. Transgenic reporters labeling both the immune cells and hepatocytes were used to count cells through the optical sections. At the juvenile and adult stages, cell counts were made on whole liver dissected and imaged *ex vivo* using a maximum intensity projection of the z-stack using the ImageJ cell counter tool. Two representative z-stacks were taken for each liver counted.

### Clodronate-mediated macrophage depletion

Transgenic larvae were injected at 3 dpf intravenously with 1 nL clodronate liposomes (Liposoma) supplemented with Alexa 568 conjugated dextran (10 kDa, Invitrogen) used at 1:100 for visualization of injection. Larvae were incubated for two days to allow macrophage depletion to occur before they were subjected to brain microinjection with LPS or water vehicle at 5 dpf. Clodronate depletion of macrophages in the brain (microglia) was confirmed in a subset of larvae using the neutral red staining assay. After brain injections, analysis of neutrophil numbers in the liver was conducted in live transgenic zebrafish imaging at 3.5 hpi.

### Whole mount RNA *in situ* hybridization

RNA *in situ* was performed using standard methods. Antisense riboprobes were synthesized from plasmids as described[72] encoding *mfap4, mpx* (Open Biosystems clone 6960294), and a 739 bp coding fragment of *irg1* (NM_001126456.1; pCES161) cloned from a cDNA library derived from a 4 dpf *E. coli* injected larva using primers Forward-5’-TCGTTCTGCCAGTAGAGATGTTA-3’ and Reverse-5’-GCGAGCTGAGATGCCTCTAAAC-3’.

### RNA Isolation, qPCR and RT-PCR

RNA was isolated following the RNAqueous-Micro kit RNA Isolation Procedure (Ambion). Whole larvae or dissected livers and remaining body were lysed in 100-300 uL RNA lysis buffer. Larval liver dissections were performed on transgenic larvae *Tg*(*lfabp10a:DsRed*) to aid in identifying the liver. cDNA was made from 150 or 200 ng of total RNA using oligo (dT) primer with SuperScript IV reverse transcriptase (Invitrogen) for qPCR or RT-PCR analysis. qPCR was performed on the QuantStudio 3 Real-Time PCR System (Applied Biosystems) using SYBR Green. The delta-delta ct method was used to determine the relative levels of mRNA expression between experimental samples and control. 1:10 and 1:4 cDNA dilutions were used for whole larvae and dissected liver tissues respectively. *ef1a* was used as the reference gene for determination of relative expression of all target genes. Primer sequences for qPCR and RT-PCR analysis are listed in Supplementary Table 1.

### Morpholino injections

Antisense morpholino oligos were purchased from Gene Tools and re-suspended in water to make 1mM or 3mM stocks. Morpholino sequences are listed in Supplementary Table 1. Morpholinos were heated at 65°C for 5 minutes and cooled to room temperature before injecting into single-cell embryos at 0.5-1 nL.

### Neutral red staining

Microglia were scored in live larvae by neutral red vital dye staining as previously described [63, 72]. In brief, 3-4 dpf larvae were stained with neutral red by immersion in fish water supplemented with 2.5 μg/mL neutral red and 0.003% PTU at 28.5°C for 1 hour, followed by 1–2 water changes, and then analyzed 2–3 hours later using a stereomicroscope.

### CRISPR-Cas9 targeted mutagenesis of *il-34*

The target gene was *il-34* (NCBI accession: NM_001128701.1; Gene ID: 560193). Co-injection of Cas9 mRNA and guide RNAs (gRNAs) was conducted in wildtype 1-cell stage zebrafish embryos. Cas9 mRNA was transcribed from XbaI linearalized pT3TS-nCas9n plasmid (Addgene #46757) using mMessage mMachine T3 Kit (Ambion) according to the manufacturer’s instructions. CRISPR targets for gRNA designs were identified using CHOPCHOP (http://chopchop.cbu.uib.no)[127]. The following gene-specific oligonucleotides using T7 promoter were used to make gRNAs as previously described[127]; gene-specific target sequences were: *il-34* gRNA-1 5’-CCGAATGCTGGCTTCTCA -3’, *il-34* gRNA-2 5’-GCTTCTCAGGGGGCTGCT -3’, *il-34* gRNA-3 5’-CCCTGAGAAGCCAGCATT-3’. Genotyping primers used: il-34 crispr F 5’-GCAGGTGTTTAATAAAGTGTG-3’ and il-34 crispr R 5’-ATCCAAAGTTCCCACATCTGAC-3’. In vitro transcription of gRNAs from assembled oligonucleotides was conducted using the HiScribe T7 Quick High Yield RNA Synthesis Kit (NEB). Injected clutches of embryos were validated to contain CRISPR mediated mutagenesis by a T7 endonuclease assay. Mutations were analyzed by TOPO TA cloning followed by Sanger sequencing.

### Small-molecule anti-inflammatory drugs

Administration of different small-molecule chemicals, DMSO control, or no treatment were performed in parallel starting at 3 dpf through the time of brain microinjection at 4 dpf in clean multi-well dishes. Analysis of the effect on macrophages was performed at 8 hpi either by RNA *in situ* hybridization or by live confocal imaging using the liver and macrophage transgenes (*fabp10a:DsRed* and *mpeg1:GFP*). 17-DMAG (5 uM) and dexamethasone (6.5 uM) were reconstituted in water and these treatments were compared to the water controls. GW2580 (25 uM), Celastrol (0.22 uM) and Bay 11-7082 (1 uM) were resuspended in DMSO so these treatments were compared to the DMSO-treated controls. List of small molecules is detailed in Supplementary Table 1.

### Liver size measurement

Whole liver was imaged 48 hpi after LPS or water injection at 4 dpf in the brain tectum on a confocal Nikon A1R+ using an apochromat lambda 40x water immersion objective (NA 1.15). 4 dpf larvae for the brain injections were derived from either uninjected, *pu*.*1*-MO injected, or negative control *p53*-MO injected wild-type embryos. Z-steps were taken at 1 um thickness. Surface and 3D rendering of the z-stack to measure the liver volume was conducted using the Imaris 3D/4D Image Analysis Software.

## Statistical analysis

Unpaired two-tailed t-tests were performed unless otherwise noted. F test was used to compare variances. For unequal variances, Welch’s correction was used on the two-tailed t-test. For multiple comparisons of 3 or more groups, one-way ANOVA test was applied followed by pairwise tests to determine the pair(s) showing significant differences. GraphPad Prism 8 was used to run statistical tests and create graphs. Scatter bar plots show symbols representing biological replicates.

## Author Contributions

LY: methodology; experimentation and formal analyses of in vivo imaging, morpholino experiments, drug tests, time-course, RNA in situ, qPCR; visualization. JAJ: methodology; experimentation and formal analyses of in vivo imaging, morpholino experiments, Kupffer cells, adult stages, and qPCR. AME: experimentation and formal analyses of CRISPR-based manipulations, liver size, and morpholino experiments; generation of new transgenic lines; contributed to original draft writing; visualization. VH and VK: experimentation and analyses of genetic mutants. CTD: technical support for RNA in situ, cDNA analysis, and imaging. CES: conceptualization; methodology; experimentation and formal analyses of brain drainage/genetic mutants/dynamic imaging; writing original draft; data analyses and visualization; supervision; funding. All authors contributed to manuscript editing.

## Acknowledgements

We thank Jonathan Gable for critical insights and discussions on our manuscript. We are grateful for the generous sharing of fish lines from Tjakko van Ham, John Rawls, David Tobin, and Brant Weinstein, and constructs from Martin Distel. We also thank the UNC Neuroscience Microscopy Core Facility for assistance with the Imaris imaging analysis and UNC Zebrafish Aquaculture Core Facility for zebrafish housing and care. L.Y. was funded by a UNC SURF fellowship, and J.A.J. was supported by a T32 ES007126 training fellowship. This work was supported by NIH NIGMS grant 1R35GM124719 to C.E.S.

