## Supplemental File for "Drainage of inflammatory macromolecules from brain to periphery targets the liver for macrophage infiltration"

### Department of Microbiology and Immunology, University of North Carolina at Chapel Hill, Chapel Hill, NC

‡ Co-lead author

§ Lead and senior author

* Corresponding author


This file contains:

Supplementary Figures 1-16 (figure legends and figures)

Supplementary Movies 1-9 (movie legends)

Supplementary Table 1 (supporting information for Materials and Methods)

Supplemental References

**Supplementary Figure 1. Whole-body analysis using RNA in situ hybridization shows abnormal localization of macrophages in the liver after brain injection of LPS or bacteria.**
**a** Schematic showing injection of substances (yellow) in the tectum parenchyma (blue) in the 4 dpf zebrafish larva and subsequent analysis of the periphery that highlights the most prominent change in the liver (red). A, anterior; P, posterior; D, dorsal; V, ventral. **b** In the 4 dpf zebrafish, macrophages (*mfap4*+) are normally absent in the liver (dotted region) while sparsely spread throughout the body (arrowheads) as shown in larvae which were uninjected or water vehicle injected in the brain at 6 hpi. By contrast, after LPS injection in the brain, strong expression of macrophage marker *mfap4* is found in the liver (dotted region), while the brain injection site is often also accompanied by a *mfap4* expression increase (arrow). **c** Analysis of zebrafish macrophage-specific markers *mfap4* and *irg1* in the liver (dotted region) after brain injections with water vehicle, *E. coli*, or LPS. Results indicate that both LPS and *E. coli* injections induce ectopic localization of macrophages in the liver, which appear to be activated (expressing *irg1*) at least in a subset, but no macrophages in the liver in control animals.

**Supplementary Figure 2. In vivo time-lapse imaging shows dynamic movements and processes of infiltrating macrophages in the liver at short- (10 hpi) and long-term (48 hpi) timepoints after brain LPS injection.**
Single slice images from z-stacks taken using a 40x objective correspond to Supplementary Movies 1 and 2. **a** Representative single-slice images of three timepoints from time-lapse imaging of macrophage infiltration at 10 hours post injection of LPS in the brain. Top, merged channels of *mpeg1:GFP* labeling macrophages and *fabp10a:DsRed* labeling hepatocytes; middle, DsRed channel showing hepatocytes and pronephros; bottom, GFP channel showing macrophages. Pronephros are labeled because the red fluorescent dextran used as a tracer for the brain injection gets into circulation and is filtered by the pronephros. **b** Representative static single-slice images of the whole liver and macrophages at 48 hpi of LPS in the brain. Yellow arrows, infiltrating macrophages; red arrow in **a**, circulating monocyte/macrophage; red arrow in **b**, macrophage with elaborate long processes intercalated between hepatocytes. Blue arrows in **b**, peripheral macrophages not in the liver.

**Supplementary Figure 3. Brain-LPS injection was not found to induce *mfap4* expression in the liver.**
**a** Comparison between control uninjected and brain-LPS injected in double transgenic 4 dpf larvae was made at 16 hpi. Fluorescent reporters for liver (*fabp10a:BFP*) and macrophages

(*mfap4:tdTomato*) were used for *in vivo* imaging to determine whether ectopic induction of *mfap4* may occur in hepatocytes due to LPS activation. Representative single 2 um z-plane images do not show ectopic expression of *mfap4* other than in infiltrating macrophages in the liver (which are stereotypically located near or within liver sinusoids or gaps between hepatocytes) after LPS injection. Coinciding with a lack of macrophage infiltration in the control animals, no *mfap4* expression was observed in the liver (demarcated by dotted line). **b** Scatter plot shows analysis of macrophage infiltration in the liver using the double transgenic larvae. Brain-LPS injection causes infiltration of *mfap4:tdTomato* expressing macrophages but not in the control uninjected animals. **c** Quantification of liver cells expressing *mfap4* shows none did. Individual slices through entire z-stack of whole liver were assessed for co-expression of hepatocyte reporter with the *mfap4* reporter. In all plots, each symbol represents an independent larva analyzed. Two-tailed student's t-test was used to determine statistical significance.

**Supplementary Figure 4. Macrophages infiltrate the liver through vasculature and vasculature-independent routes.**

Double transgenic zebrafish expressing the macrophage *mpeg1:GFP* and endothelial *kdrl:mCherry* reporters were used to localize infiltrating macrophages after brain-LPS injection at 8-10 hpi in the 4 dpf zebrafish. **a-f** Representative images corresponding to Supplementary Movie 3. **a, b, c** 3D volumetric view of whole liver (demarcated by a dotted line) and surrounding region taken from three timepoints of a confocal time-lapse imaging (see Supplementary Movie 3) of an 80 um z-stack at 40x. **d, e, f** High magnification of the boxed region shown in the corresponding left panels. Representative infiltrating macrophages are labeled numerically 1-5 with their relative position to the vasculature: inside (white arrow), associated (blue arrow), or independent of vasculature (yellow arrow).

**Supplementary Figure 5. Long-term in vivo tracking of infiltrating macrophages shows that occupation time in the liver can be used for their classification.**

**a** Representative 3D images from 6 timepoints corresponding to Supplementary Movie 4 from a 4 dpf zebrafish which was brain LPS injected and imaged starting at 12 hpi. Infiltrating macrophages can be classified into three groups: circulating (present in only a single timepoint), transient ($\leq$ 2 hours) and residing cell (> 2 hours). Double transgenic zebrafish labeling macrophages (*mpeg1:GFP*) and hepatocytes (*fabp10a:DsRed*) were used to track macrophage infiltration over a 10-hour period. Color overlay of both channels and single GFP channel in grayscale for macrophages are shown. LPS was co-injected with Alexa 568 conjugated dextran to validate brain injections and detect systemic distribution of the injected material based on labeling of the pronephros by the fluorescent dextran. Labeling of pronephros has been previously described after intravenous injection of a fluorescent tracer[1], but labeling of proximal kidney tubules can also be observed in transgenic lines expressing a red fluorescent protein. **b** 2D projection of the tracking of infiltrating macrophages in the entire liver over all timepoints of analysis. Each traced cell is represented by a circle for its location at each timepoint if present in the liver, and a line for its movement between the timepoints. Different colors represent different cells. **c** Relative percentage of the different types of infiltrating macrophages based on liver occupation. n= 47 macrophages were tracked and analyzed. **d** Scatter plot shows the total duration of each traced infiltrating macrophage in the "transient" and "residing" groups. **e** Total distance and average speed of each macrophage in the "transient" and "residing" groups. Total distance was not significantly different, but the "transient" macrophages moved faster with a significantly higher average speed than the "residing" cells. sd, standard deviation; ns, not significant.

**Supplementary Figure 6. Injected LPS macromolecules disperse from the hindbrain ventricle into the hindbrain and spinal cord/trunk interstitial spaces and are localized within the head and trunk lymphatics.**

Wild type double transgenic zebrafish at 4 dpf were microinjected with fluorescently tagged LPS in the tectal brain. To localize the LPS relative to the lymphatic and vasculature structures, two transgenes were used: *mrc1a:GFP (cite)* and *kdrl:mCherry (cite)*, respectively. Since *mrc1a:GFP* is known to also be expressed by macrophages (cite), we generated a new macrophage reporter (*mpeg1:BFP*) to provide a third channel in blue to distinguish lymphatic from macrophage expressions. Blue arrows point to microglia/macrophage. **a** Three multi-tiled representative frontal plane optical slices from an *in vivo* confocal z-stack at 1.5 hpi show top, middle, and bottom sections through the head and anterior trunk of the LPS-injected animal. **b** Schematic of the frontal plane optical sections taken by *in vivo* multi-tiled z-stack confocal imaging. **c, d** High magnification images corresponding to dotted region in **a**. Most superficial layer of the CNS where LPS (red) is concentrated in the hindbrain ventricle and adjacent interstitial parenchymal space (c, asterisk) as well as the most-dorsal junction of the hindbrain-spinal cord (d, asterisk). **e, f** High magnification images of dotted region in middle slice shown in **a**. LPS is found localized to the lymphatic structures (blec and islv), and abundant in the interstitial space extending from the hindbrain ventricle (asterisk). **g,h** High magnification images corresponding to bottom slice in **a**. LPS is highly abundant in the spinal canal (sc) as well as tectal and hindbrain lymphatic cells (blec). **i** Top and middle slices of a different LPS-injected animal show the same distribution of LPS at 1.5 hpi. LPS does not appear to localize within the blood vasculature in the CNS or trunk to the same extent as it overlaps with the lymphatic cells. Top slice shows vasculature DLAV surrounded by but not overlap with LPS, arrows. *, interstitial space; DLAV, dorsal longitudinal anastomotic vessels; DLLV, dorsal longitudinal lymphatic vessel; blec, brain lymphatic endothelial cells; islv, intersomitic lymphatic vessels; sc, spinal canal; hv, hindbrain ventricle; hb, hindbrain; lym, lymphatics. Skin is highly auto-fluorescent and labeled by all channels at the border of every tissue. Scale bars all show 50 um.

**Supplementary Figure 7. Injected LPS macromolecules into tectal brain results in distribution of LPS within the facial lymphatic network and peripheral vasculature.**
**a** Schematic of the sagittal optical planes taken by *in vivo* multi-tiled z-stack confocal imaging. **b** Whole-mount side view of a representative LPS-injected 4 dpf larvae carrying the vasculature reporter *kdrl:mCherry* at 12 hpi showing LPS strongly retained in the hindbrain ventricle (hv) and surrounding brain interstitial space as well as the brain and facial lymphatics (blec and FL, arrows) and the peripheral vasculature (PCV and PHS, arrows), but not in the brain blood vessels (vas). **c** High magnification of dotted region in **b** showing uptake of LPS as fluorescent puncta (arrows) by the PCV, but LPS is absent in the hepatic region. **d** Characterization of the same timepoint at 12 hpi as in **b-c** using fish carrying the lymphatic reporter clearly show localization of LPS within the facial lymphatics (FL, arrows). By stark contrast, LPS is not localized to the brain vessels, thereby providing evidence for drainage of LPS from brain to peripheral circulation through the lymphatics via the interstitial fluid. FL, facial lymphatics; PHS, primary head sinus; PCV, posterior cardinal vein; blec, brain lymphatic endothelial cells; sc, spinal canal; hv, hindbrain ventricle; vas, brain vessels; mb, midbrain; *, interstitial space. Scale bars all show 50 um.

**Supplementary Figure 8. Microinjection of fluorescent LPS into brain tectum at 4 dpf results in low LPS transfer from hindbrain ventricle into brain interstitial space at around 30% and in the periphery at less than 15% of injected LPS level.**
**a** Time plot showing LPS level changes in different regions as a ratio of injected fluorescent LPS. Injected LPS level was determined by the relative saturation level of fluorescence in the hindbrain ventricle as it is the immediate site filled in by LPS at time of injection. Both hindbrain and telencephalon ventricles reach 100% maximum LPS level based on fluorescence, while hindbrain interstitial space reaches around 30% of maximum and the periphery using accumulation in the pronephros as a proxy reaches less than 15% of maximum. This plot illustrates a small fraction

of maximum LPS that does get passed onto the periphery. Plot shows average and s.e.m. (standard error of means) of data from three independent LPS-injected animals. See Figure 2 for related analysis of tracing LPS from brain microinjection. **b** Still images of key events from tracing LPS-Alexa 594 after brain injection by high-resolution time-lapse stereomicroscopy, taken from a different LPS-injected animal than the example shown in Figure 2. Injection occurred at the 4s timepoint. Some injected animals such as this one, LPS is detected to also move anteriorly from midbrain/hindbrain ventricle into telencephalon ventricle. By 48 minutes post injection, we detect broad distribution of LPS in circulation at a low intensity. FL, facial lymphatics; blec, brain lymphatic endothelial cells; hbv, hindbrain ventricle; mb, midbrain; hb, hindbrain; tele, telencephalon; m, minute; s, second.

**Supplementary Figure 9. Blocking circulation by a morpholino-mediated tnnt2 knockdown prevented macrophage infiltration into the liver 8 hours after brain injection with LPS.**
**a** Representative 3D volumetric and single z-plane images shown for after LPS or water vehicle injection in the brain. Dotted region shows liver in the single channel panels. Transgenes *fabp10a:DsRed* and *mpeg1:GFP* were used to label hepatocytes and macrophages, respectively. **b** Quantification of the number of macrophages found in the liver at 8hpi in 4 dpf zebrafish. No difference was found between the control and LPS injected groups as they both lacked macrophage infiltration. Each symbol represents an independent animal. Statistical test was determined by a two-tailed t-test. ns, not significant; MO, morpholino.

**Supplementary Figure 10. A test of anti-inflammatory drugs revealed dexamethasone and Bay 11-7082 to be effective for preventing liver infiltration by macrophages.**
**a** Frequency of liver infiltration at 8-10 hours after brain-LPS injection showed that only Bay 11-7082 (1 uM) and dexamethasone (6.5 uM) treatments were effective for preventing macrophages from infiltrating the liver as determined by in situ hybridization. Number of independent animals as shown in the parenthesis. **b** Representative whole-mount RNA hybridization for macrophage marker *mfap4* showing liver in the dotted region. White dotted liver indicates infiltration, while black dotted liver shows no infiltration. **c** Quantification of the number of infiltrating macrophages either after water or LPS injection in the brain in the control or drug treated conditions as determined by in vivo imaging. **d** Representative single z-plane images from z-stacks that show abundant infiltrating macrophages (arrows) in control treated animals after brain-LPS injection, but no infiltration in dexamethasone or Bay 11-7082 treated larvae. Student's t-test was used to determine statistical significance.

**Supplementary Figure 11. Characterization of indel mutations created in Cas9/*il-34* gRNAs injected *il-34* crispant animals confirms efficacy of disrupting *il-34* coding sequence.**
**a** Representative sequences of indel mutations from three transient *il-34* crispant animals all showing frameshift mutations leading to nonsense proteins often with an early termination (red boxes highlight corresponding translation). The exon containing the *il-34* start codon (yellow box), three targeting gRNAs (teal boxes) and the *il-34* reverse sequencing primer (green box) are shown. Blue boxes show number of nucleotide change with a "+" for insertion and a "–" for deletion (del). Top bar shows consensus in the alignment: green for complete match, yellow and red indicate some mismatches. **b** Relative frequency of different indel mutations based on net nucleotide change from the wildtype sequence. Each distinct mutation was counted.

**Supplementary Figure 12. Hepatic response to brain stimulation still evident after establishment of Kupffer cells and adult vasculature/brain structures in juvenile adults.**
**a** Live *ex vivo* imaging of dissected liver from juvenile adult zebrafish after microinjection of either water vehicle or LPS in the brain at 16-18 hpi. Top, arrows point to macrophages in liver using the *mpeg1:GFP* transgene. Bottom, arrows point to neutrophils in the liver using the *lyz:mCherry* transgene. **b** Quantification of macrophage density (# *mpeg1:GFP*+ cell per field of view, FOV). **c** Quantification of neutrophil density (# *lyz:mCherry*+ cell per FOV). Scatter plots show uninjected controls, or animals microinjected with water vehicle or LPS in the brain. Each FOV equals 0.045 squared mm. Statistical significance was determined by a two-tailed Student's t-test.

**Supplementary Figure 13. Clodronate-mediated macrophage depletion results in an increase in neutrophil infiltration after brain-LPS injection.**
**a** Schematic illustrating the method. Intravenous injection of clodronate liposome was performed at 3 dpf to allow time for the clodronate to effectively induce cell death to most macrophages over a 48-hour period. **b** Assessment of macrophage depletion by neutral red staining shows that most animals injected with clodronate liposome indeed resulted in an apparent loss of macrophages in the brain (58%, n=12). Arrow indicates microglia in the "no depletion" category. **c** Scatter plot shows number of neutrophils infiltrating the liver to be even more increased after macrophage ablation in response to brain-LPS injection. Since clodronate based macrophage depletion is effective only in a proportion of the animals injected, we also used the cluster of the largest numbers of infiltrating neutrophils as a subset to show the animals with the most significant increase. **d** 3D volumetric view of the whole liver from imaging transgenic reporters

*fabp10a:DsRed* for hepatocytes and *lyz:GFP* for neutrophils. Left, merged channels. Middle, DsRed channel. Right, GFP channel. Brain-water sample is shown 60 um beneath the first surface of the liver, and the brain-LPS sample is 45 um below the most outer liver surface. Significantly higher number of neutrophils (arrows) is easily visualized in the brain-LPS sample. Dextran-Alexa 488 was co-injected into the brain as a tracer to validate injections, and can be seen labeling the pronephros in the GFP channel. Statistical significance was determined by a two-tailed t-test and with Welch's correction for unequal variances as determined by a F-test.

**Supplementary Figure 14. CSF3R/GCSFR knockdown was effective in reducing neutrophils for revealing neutrophil effects on macrophages.**

**a** *csf3r* morphants have less neutrophils (arrows) compared with controls as assessed by RNA in situ hybridization for a neutrophil marker myeloid-specific peroxidase mpx. Right bar graph, quantification of the area of mpx expression in the body posterior to the head using ImageJ show an average of a 27% reduction. Statistical significance was determined by a two-tailed t-test with Welch's correction. **b** Reduction of neutrophils leads to a decrease in macrophage infiltration as assessed by RNA in situ hybridization as an alternative approach in addition to live transgenic imaging as shown in Figure 4. As a control, we also verified the level of neutrophil reduction in *csf3r* morphants by *mpx* in situ. Right bar graphs show frequency of liver infiltration by each myeloid cell type. N is shown in parenthesis and represents number of independent animals analyzed. **c** Left, scatter plot representing the effects of *csf3r* morpholino-mediated knockdown on macrophages and neutrophils showed no significant change to overall macrophage numbers, but a significant reduction in neutrophils in *csf3r* morphants. *In vivo* confocal imaging of double transgenic zebrafish carrying the macrophage reporter (*mpeg1:GFP*) and the neutrophil reporter (*lyz:mCherry*) was conducted. Right, representative fluorescent maximum projection, multi-tiled images of the transgenic zebrafish body. Dotted region shows area of cell number quantification. Statistical significance was determined by a one-tailed t-test. Each symbol represents an individual larva. Scale bar represents 500 um. ns, not significant.

**Supplementary Figure 15. Serum amyloid A (*saa*), a major acute phase response gene, is not required for liver infiltration by macrophages after brain-LPS activation.**

Scattered dot and bar plot shows no significant difference in number of macrophage infiltration in the liver between homozygous *saa*$^{rdu60}$ mutants compared with their wild-type and heterozygous siblings after 16-24 hours post brain-LPS injection at 4 dpf. As a negative control, brain-water injected siblings at 16-24 hpi in the 4 dpf zebrafish did not show liver infiltration by macrophages as expected. Each circle represents an individual larva analyzed. Statistical significance was determined by a two-tailed t-test.

**Supplementary Figure 16. Depleting myeloid leukocytes to prevent immune cell infiltration in liver leads to an increase in liver growth during an inflammatory response.**

**a** Scattered dot and bar plot shows after brain-LPS injection, a relative liver size decrease in baseline and in control-*p53*-MO injected wild-type animals, whereas a significant increase in liver size in myeloid-depleted *pu.1* morphants. Relative liver size was calculated based on changes in volume of whole liver relative to the water-injected controls. Each dot represents an individual larva analyzed. **b** Representative 3D rendered volumetric images of the whole liver from all groups and conditions analyzed. Statistical significance was determined by a two-tailed t-test for pair-wise comparison, and one-way ANOVA with Brown-Forsythe correction for the three-way comparisons. All scale bars represent 50 um.

**Supplementary Movie 1. Time-lapse imaging of macrophages infiltrating the liver 10 hours after brain LPS injection.**

Representative single z-plane through the liver (*fabp10a:DsRed+*) shown from confocal imaging of one z-stack every 1 minute and 15 seconds for ~ 1 hour using a 40x objective. A range of dynamic macrophage (*mpeg1:GFP+*) behaviors is shown: some nestled in gaps between hepatocytes presumably in the sinusoids while others either circulate or traverse the liver back and forth with long processes. Left panel shows the merge channel and the right panel shows single GFP channel for macrophages. Movie file shown at 30 fps. See Supplementary Figure 4 for additional description. Arrows, infiltrated macrophages. Dotted line, liver area.

**Supplementary Movie 2. Time-lapse imaging of macrophages infiltrating the liver 48 hours after brain LPS injection.**

Representative single z-plane through the liver (*fabp10a:DsRed+*) shown from confocal imaging of one z-stack every 1 minute for ~ 1 hour using a 40x objective. Macrophages (*mpeg1:GFP+*) in the liver (*fabp10a:DsRed+*) appear to be more stationary than at the earlier timepoint at 8 hpi. Varied morphology still apparent from individual macrophages with long processes to a rounded cell shape with little to no apparent processes. Movie file shown at 30 fps. See Supplementary Figure 4 for additional description.

**Supplementary Movie 3. Association of infiltrating macrophages with the vasculature in vivo 8 hours after brain LPS injection.**

3D view of a time-lapse imaging of an 80-um volume from 4 dpf zebrafish injected with LPS showing the liver region encompassing the hepatic vasculature (*kdrl:mCherry+*) and macrophages (*mpeg1:GFP+*). One z-stack was acquired every 90 seconds for over a 5-hour period using a 40x objective. Three types of macrophage association with vasculature observed: inside, associated, or independent of vasculature. See Supplementary Figure 2 for additional description.

**Supplementary Movie 4. In vivo long-term tracking of infiltrating macrophages in the liver at 12 hours after brain LPS injection for a 10-hour continuous period.**

3D view of a time-lapse imaging corresponding to a 6-um volume collected every 2 minutes for a 10-hour period starting at 12 hpi at 4 dpf using a 40x objective. Left panel, merged channel for hepatocytes (*fabp10a:DsRed*) and macrophages (*mpeg1:GFP*). Right panel, GFP channel alone for showing macrophages. Dextran-Alexa 568 was co-injected into the brain as a tracer to validate injections, and can be seen labeling the pronephros in the DsRed channel. Circulating, transient, and residing macrophages can be observed within the liver tissue with varied cellular dynamics from being rounded and rapidly flowing through the liver to being ramified and migrating back and forth traversing the liver, respectively. See Supplementary Figure 3 for additional description. Movie file shown at 30 fps.

**Supplementary Movie 5. Time-lapse imaging of whole-body response to LPS microinjection in the brain shows recruitment of macrophages to the liver.**

3D images of whole-body from time-lapse imaging of 4 x 200 um z-stack tiles stitched into one large frame taken at 5 um z-steps using a Plan Apo lambda 20x objective every 10 minutes over a 24-hour total period starting at ~15-20 minutes post brain-LPS injection at 4 dpf. A sub-volume of the whole stack is shown from 60 um beneath the most exterior body surface in order to remove obstructive tissue layers blocking the view of the liver. Macrophages are shown by the *mpeg1:GFP* reporter, LPS-Alexa 594 is labeled by red fluorescence, and dextran is visualized by its cascade blue fluorescent tag. Macrophages throughout the body are highly mobile upon brain perturbation, but over time, a number of these cells is found to be restricted inside the liver in contrast to their fast movement through most of other organs and tissues. Fast moving macrophages throughout body were found to mostly express bright *mpeg1:GFP* as opposed to the moderately weaker reporter expression by liver-infiltrating macrophages. See Figure 2 for additional description of this imaging analysis. Movie file shown at 30 fps.

**Supplementary Movie 6. Video showing the initial restriction of brain microinjection to the brain parenchyma, ventricles, and spinal canal using a fluorescent dextran tracer in the 4 dpf zebrafish.**

Live recording of the brain microinjection of Alexa 568 conjugated dextran (10 kDa) at faster than video rate (> 30 frames per second, fps) using a Leica M165 FC stereomicroscope with a high speed and high sensitivity deep-cooled sCMOS camera (DFC9000 GT). Video shows the left side profile of a live wildtype 4 dpf zebrafish prior to injection using brightfield imaging followed by the fluorescent dextran injection as shown by the overlay of epi-fluorescence with the brightfield. The fine capillary needle is shown penetrating the injection site in the left brain tectum of the zebrafish. Immediately after injection, the injected substance rapidly fills the brain ventricles that contain the cerebrospinal fluid, and subsequently the central canal of the spinal cord. Video represents a total of 5 seconds in real time.

**Supplementary Movie 7. Rapid *in vivo* tracking of LPS movement in real time starting before brain tectal injection to nearly 1 hour post injection.**

Representative live recording of brain microinjection of LPS-Alexa 594 and its immediate aftermath in a 4 dpf zebrafish larvae at a high temporal resolution at 1 frame per second (fps) on a Leica M165 FC stereomicroscope with a high speed sCMOS camera (DFC9000 GT). LPS were found to concentrate in the hindbrain ventricle (hbv) immediately after injection and disperse along the ventricular system including the spinal canal. From the hbv, a low level of LPS were exuded out from the posterior end (arrowheads) as well as from the top arms (arrowhead) into the hindbrain interstitial space. Appearance of LPS, albeit weak, can be detected along the facial lymphatics (FL) as well as in the peripheral circulation as represented by the accumulation in the pronephros. Over time, brain lymphatic endothelial cells (blec) were found to accumulate LPS from the brain interstitial fluid. See Figure 2 and Supplementary Fig. 8 for the kinetic analysis of the LPS tracing, and Supplementary Figs. 6 and 7 for high cellular resolution analysis of LPS localization. Movie file represents a total of 50 minutes and 41 seconds of tracking, shown at 300 fps (300x faster than original process).

**Supplementary Movie 8. Dynamic macrophage-neutrophil interactions following brain LPS microinjection.**

Confocal time-lapse imaging of a 33 um z-stack was performed in the region surrounding the liver in double transgenic zebrafish carrying the macrophage (*mpeg1:GFP*) and neutrophil (*lyz:mCherry*) reporters at 4 dpf. Individual macrophages (GFP+) are prevalent around the liver but has not infiltrated the liver at 3 hpi. By contrast, a few neutrophils (mCherry+) were seen to enter liver, often passing through. A 40x objective was used to acquire a z-stack every 2 minutes for a total 1-hour period. Movie file shown at 30 fps. See Figure 4l for additional description.

**Supplementary Movie 9. Dynamic macrophage-neutrophil interactions normally surrounding the liver.**

Confocal time-lapse imaging of a 33 um z-stack was performed in the region surrounding the liver in double transgenic zebrafish carrying the macrophage (*mpeg1:GFP*) and neutrophil (*lyz:mCherry*) reporters at 4 dpf. Individual macrophages (GFP+) and neutrophils (mCherry+) are readily found to intermingle intimately as shown in this video representative. A 40x objective was used to acquire a z-stack every 2 minutes for a total 1-hour period. Movie file shown at 30 fps. See Figure 4k for additional description.

**Supplementary Figure 1.**

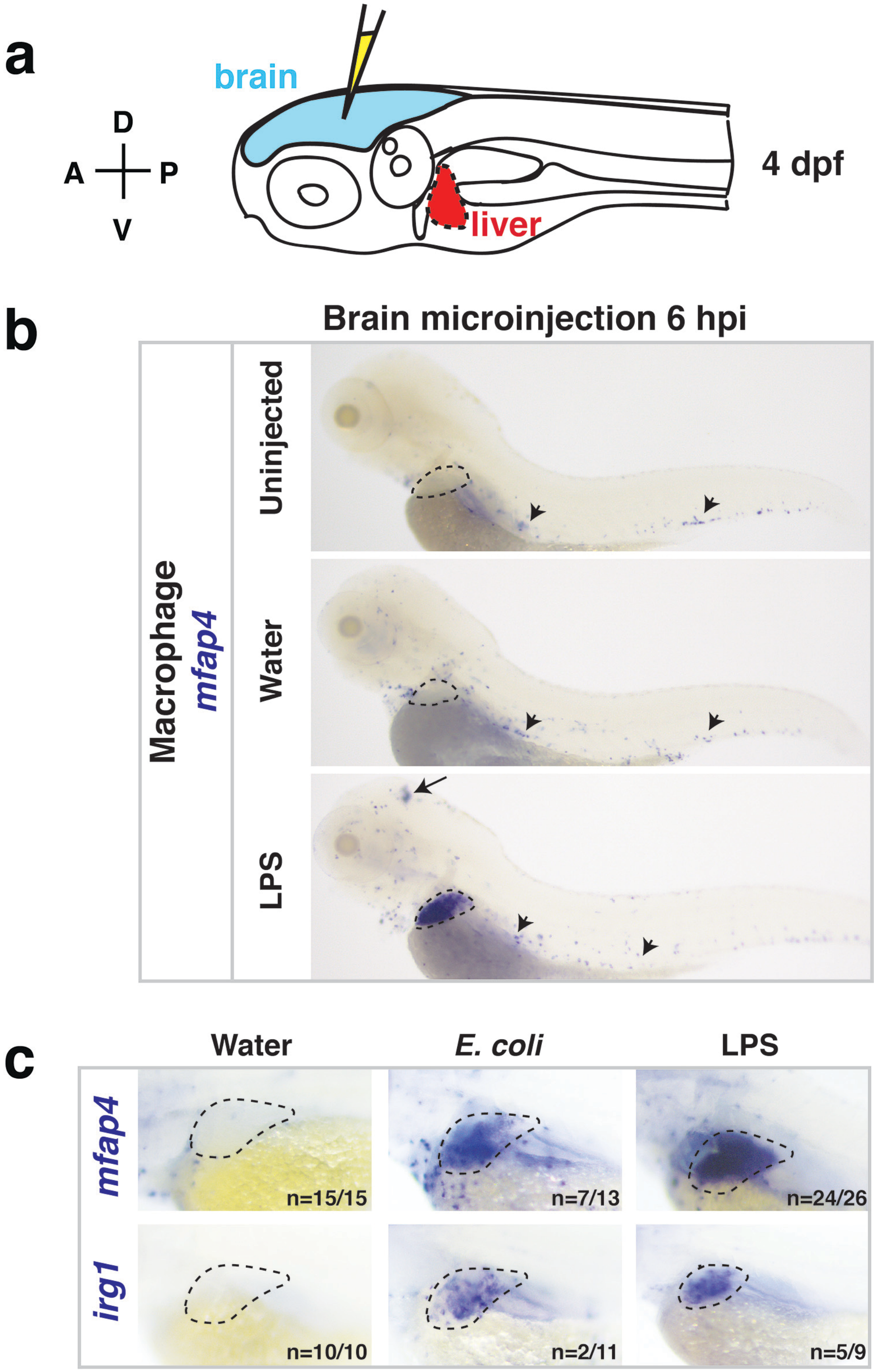

**Supplementary Figure 2.**

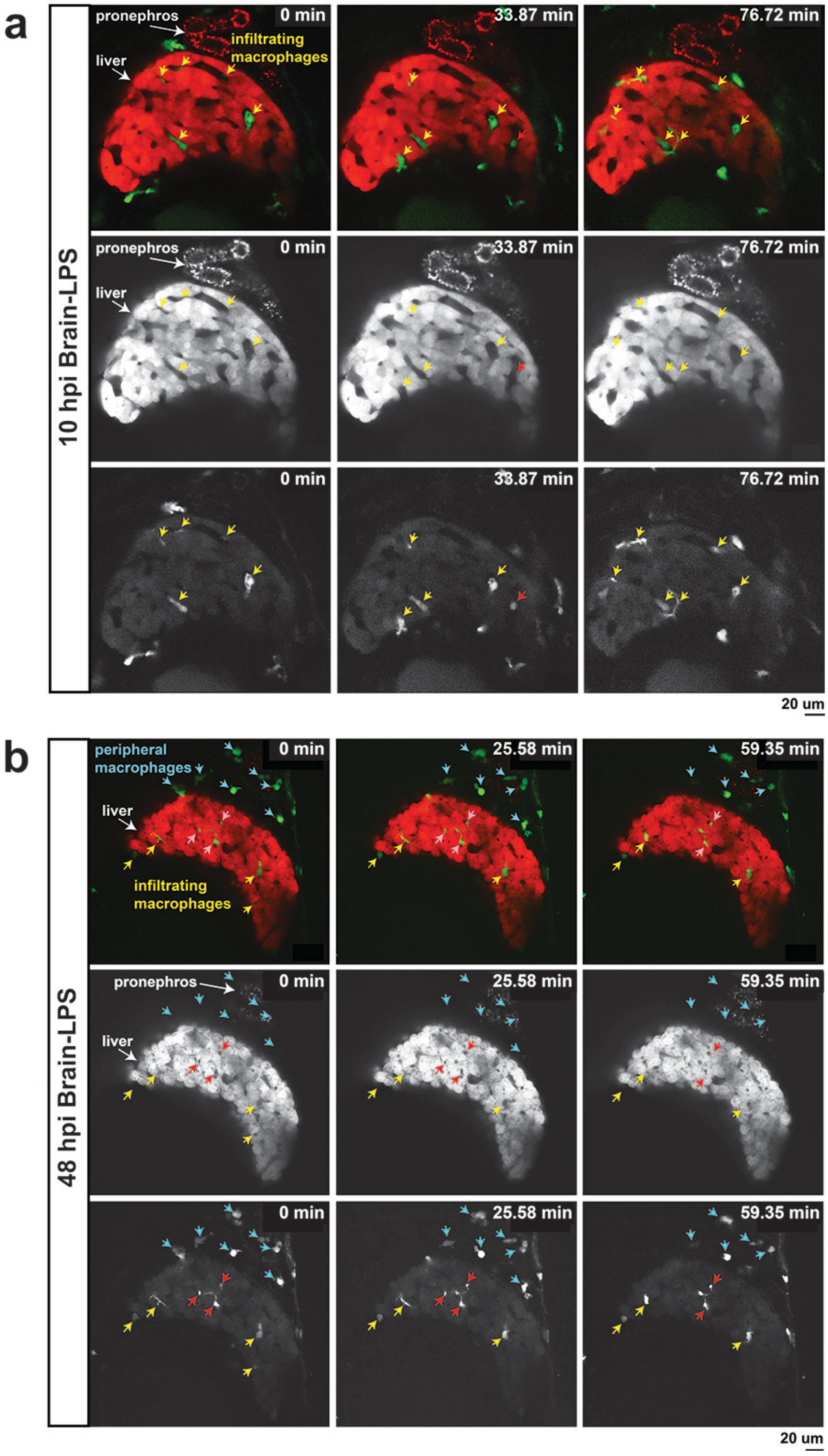

**Supplementary Figure 3.**

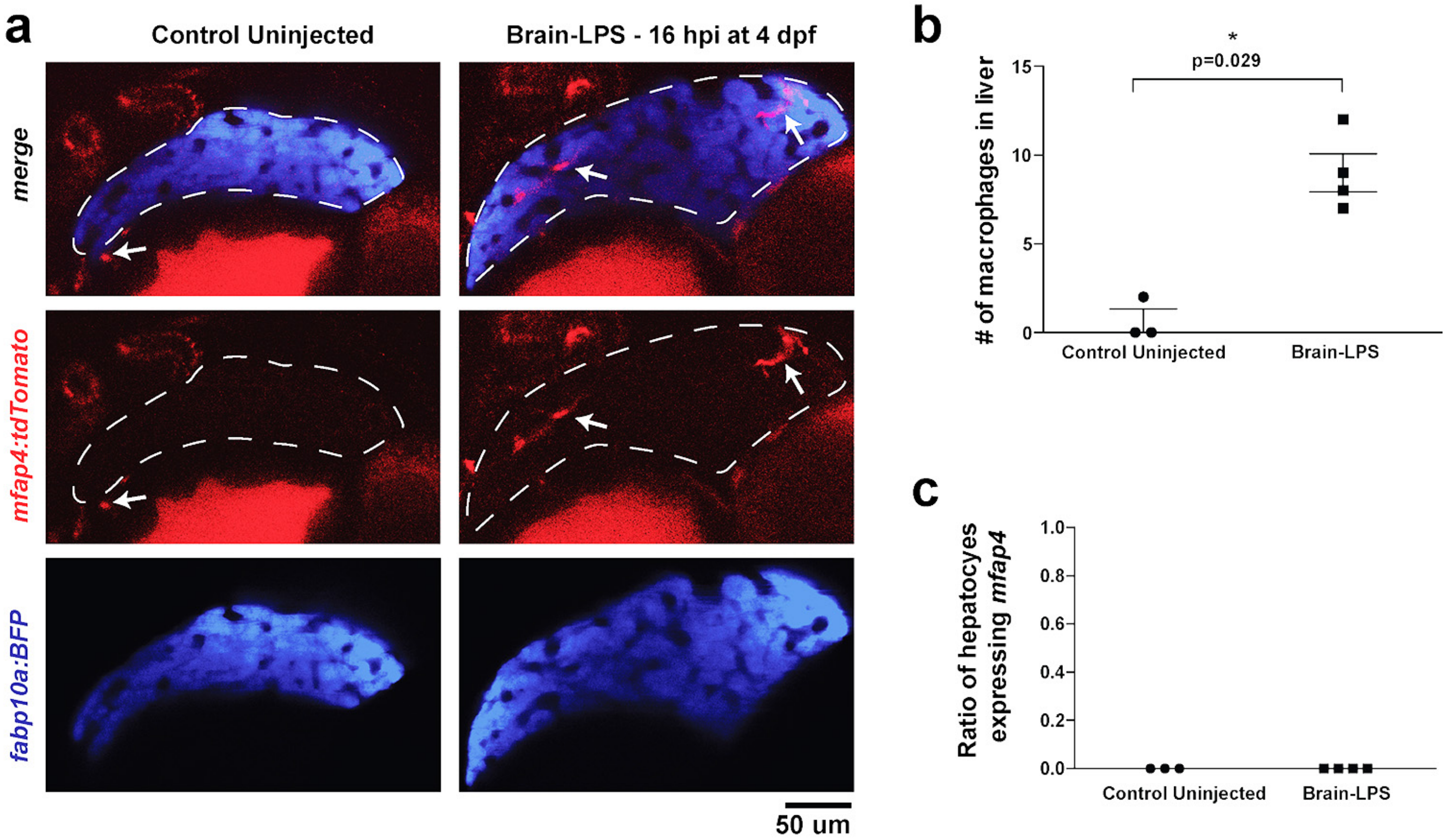

**Supplementary Figure 4.**

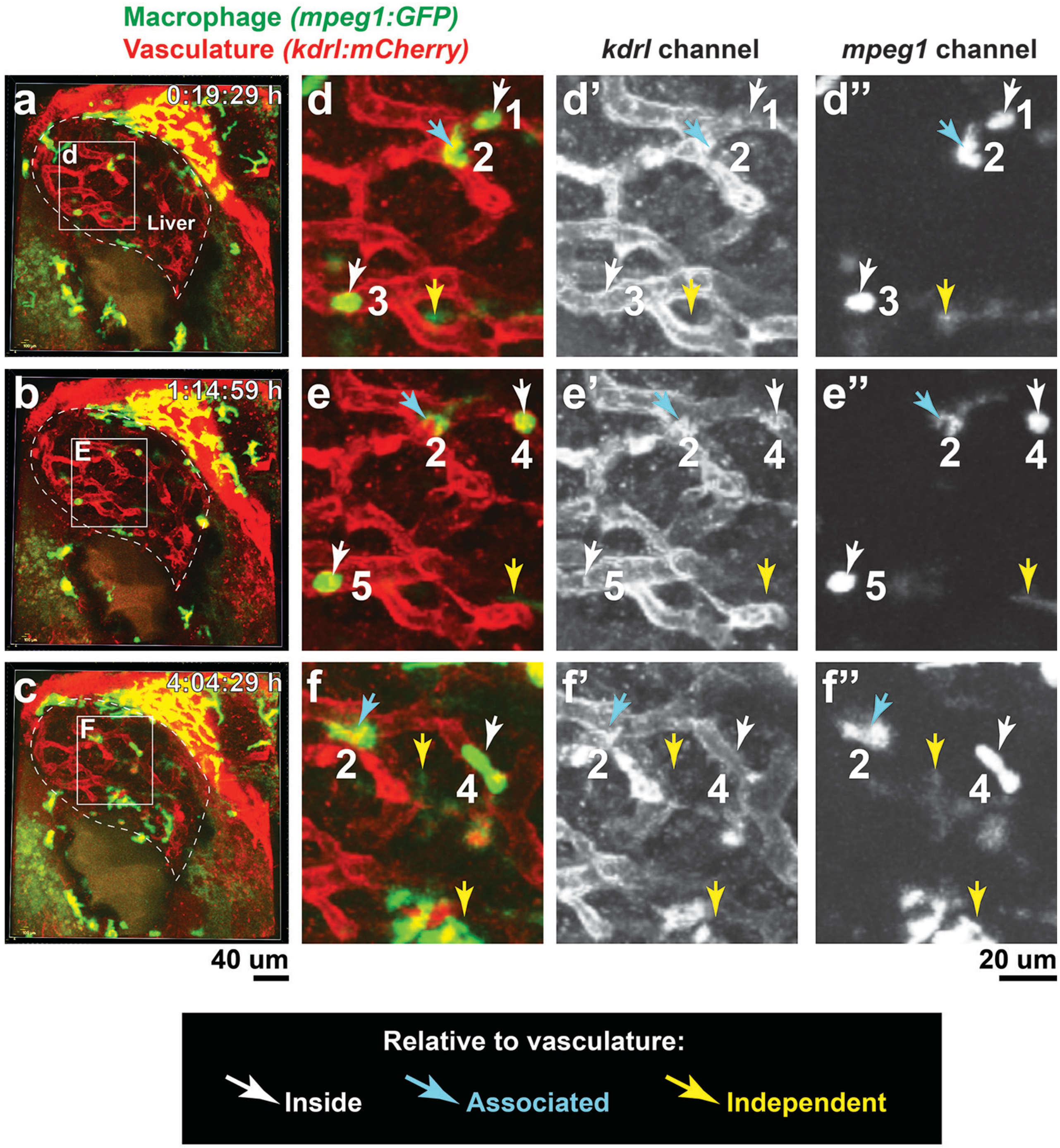

**Supplementary Figure 5.**

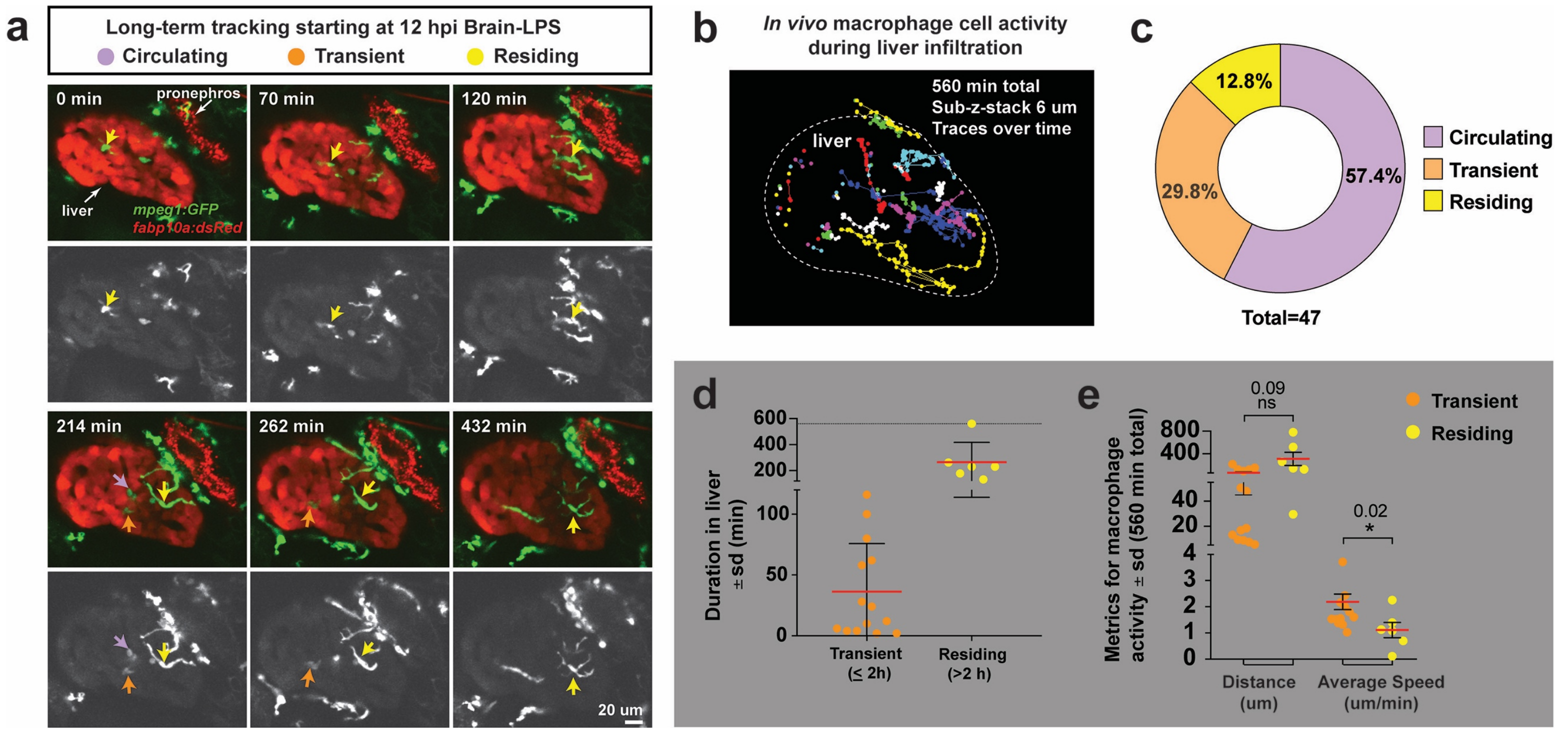

**Supplementary Figure 6.**

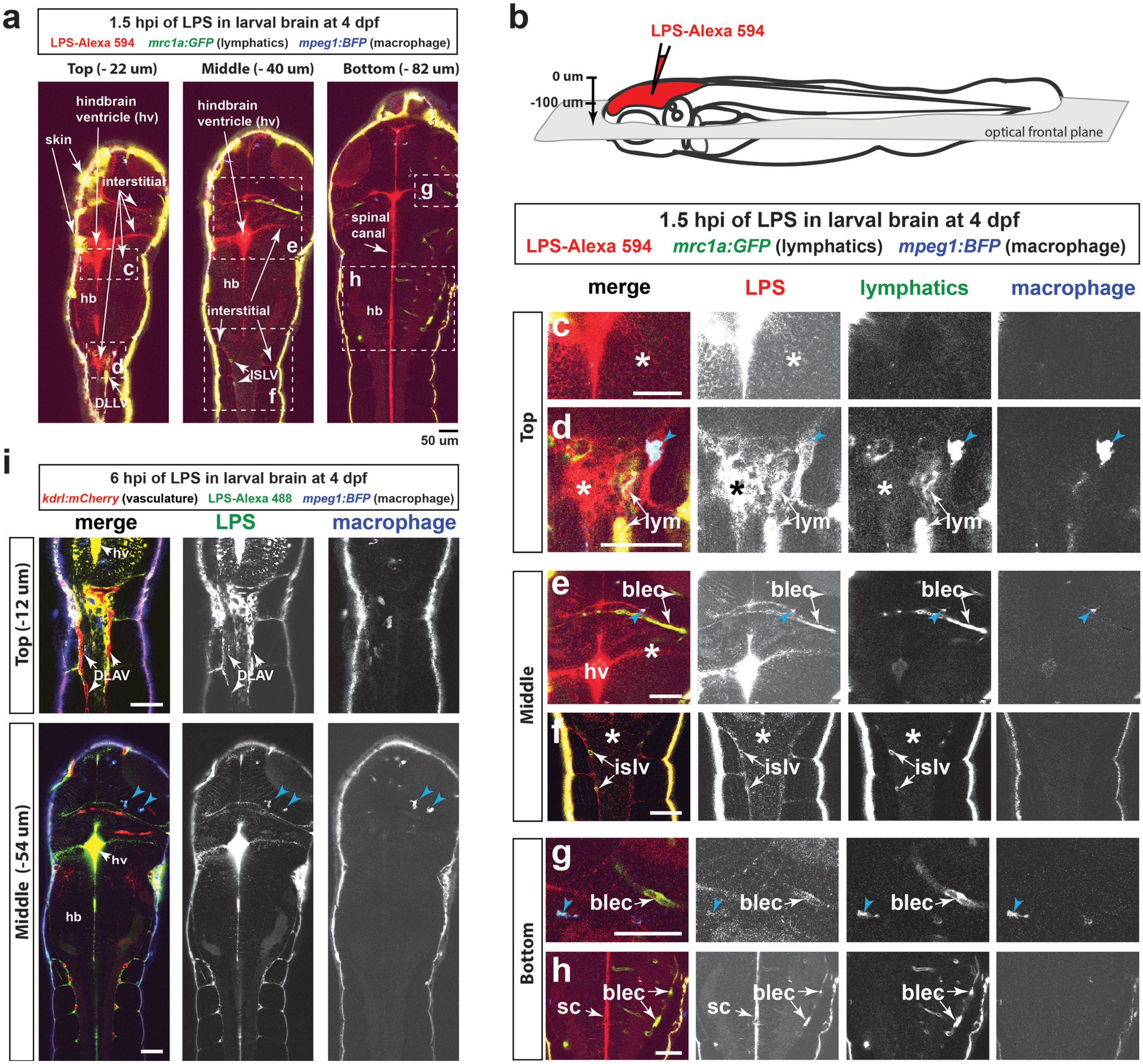

**Supplementary Figure 7.**

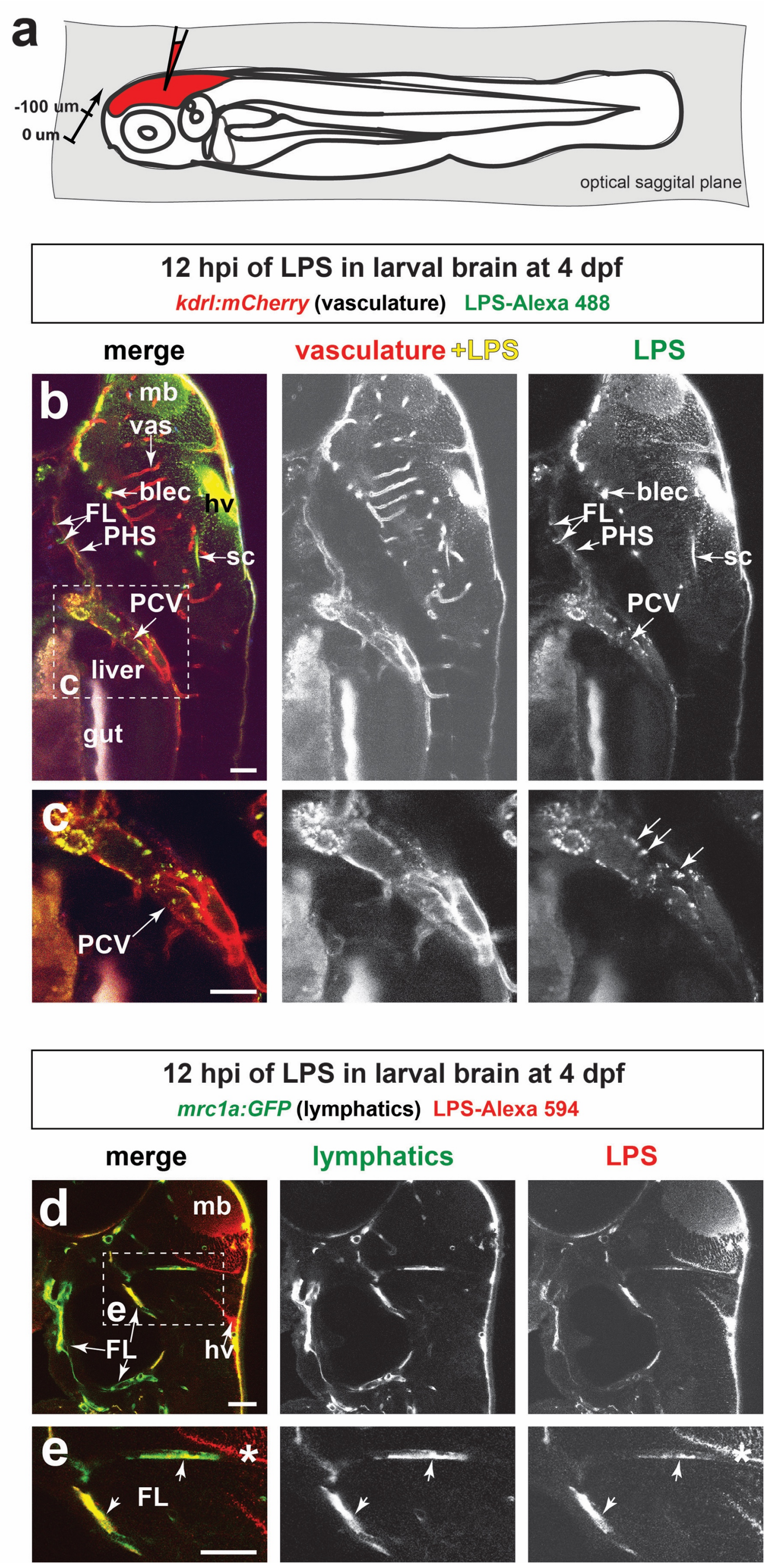

**Supplementary Figure 8.**

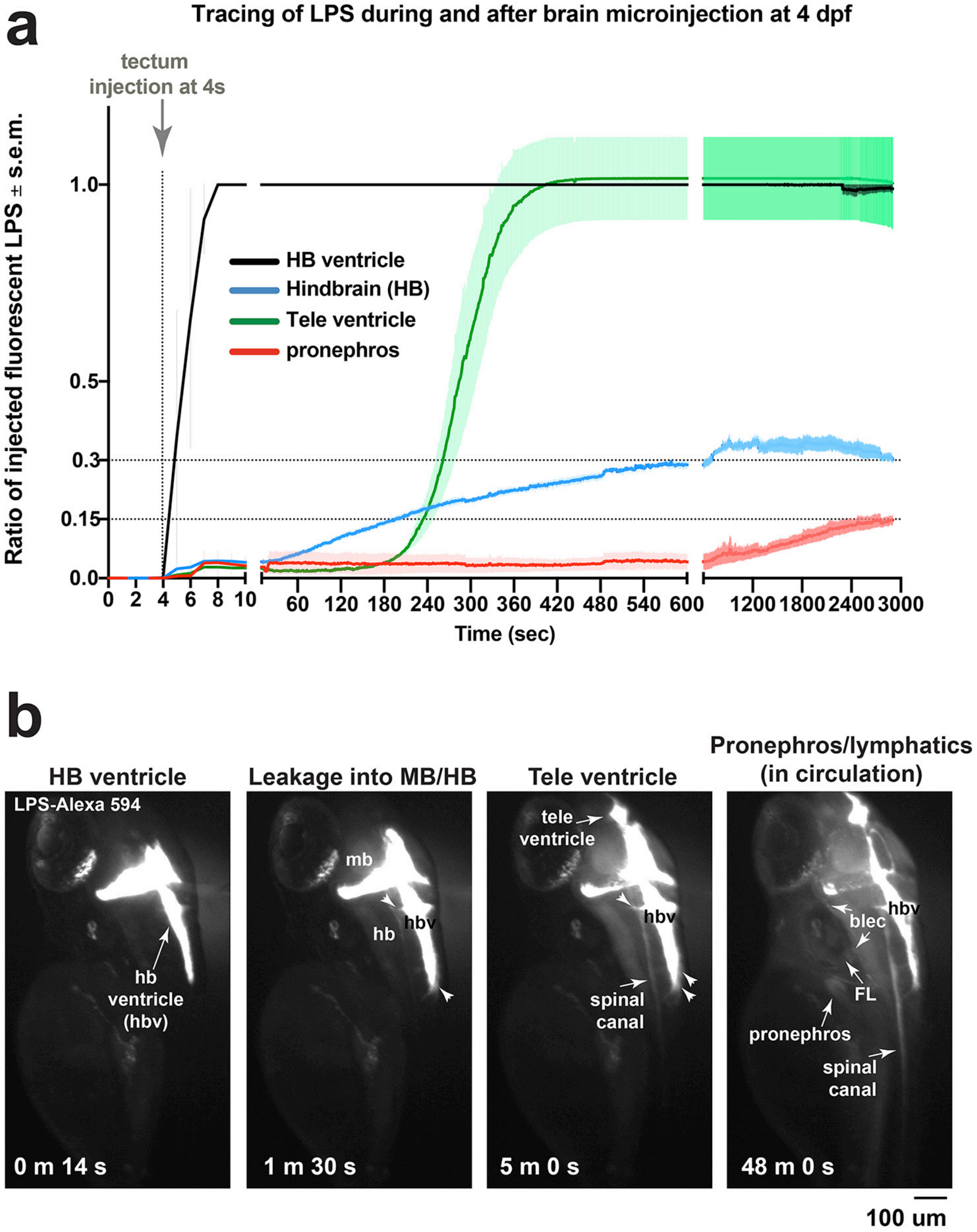

**Supplementary Figure 9.**

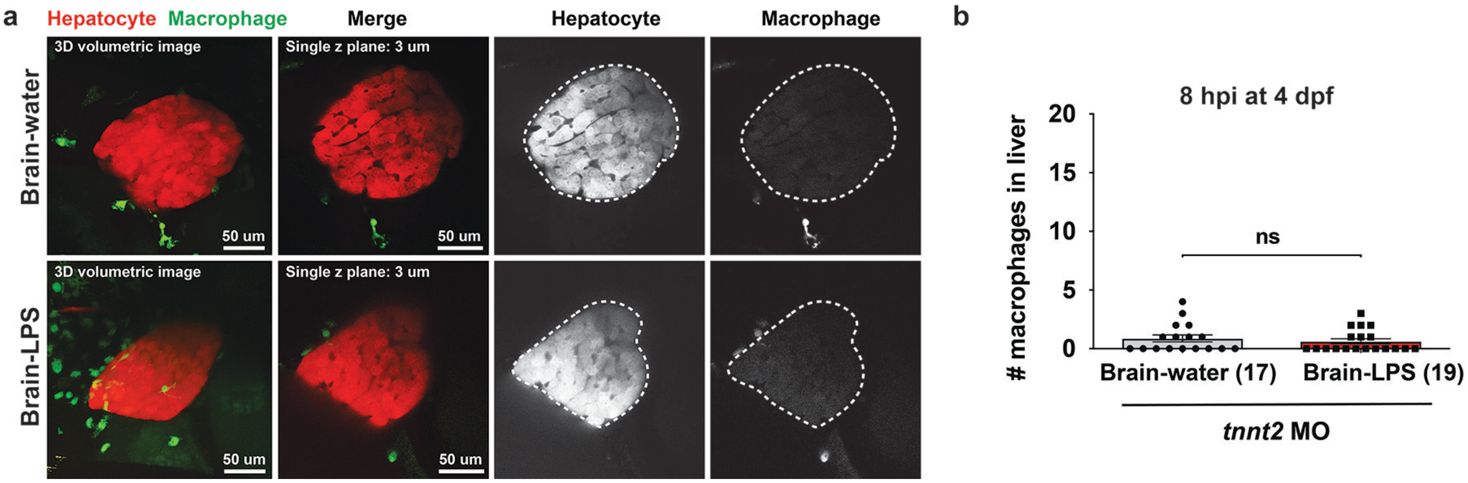

**Supplementary Figure 10.**

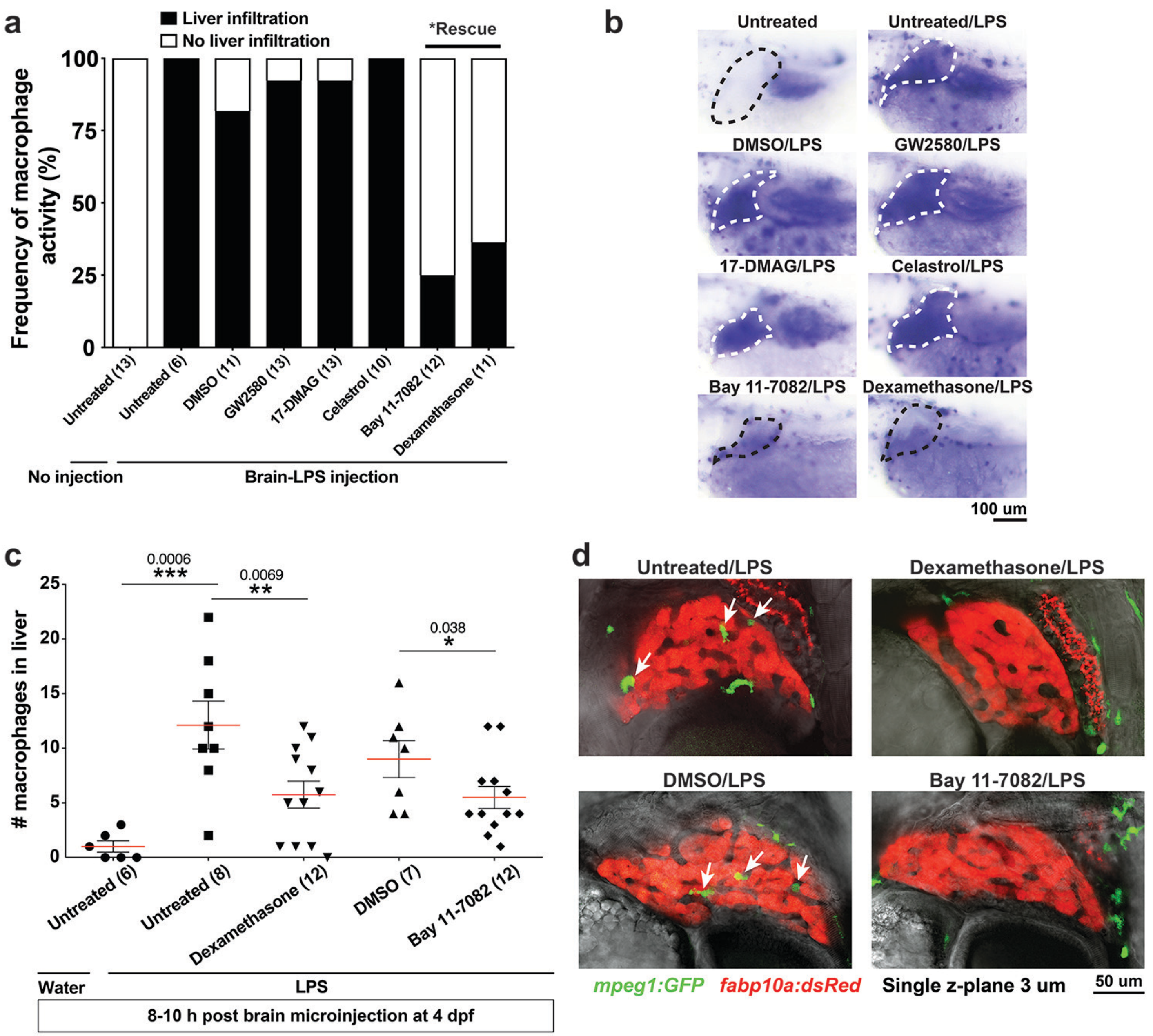

**Supplementary Figure 11.**

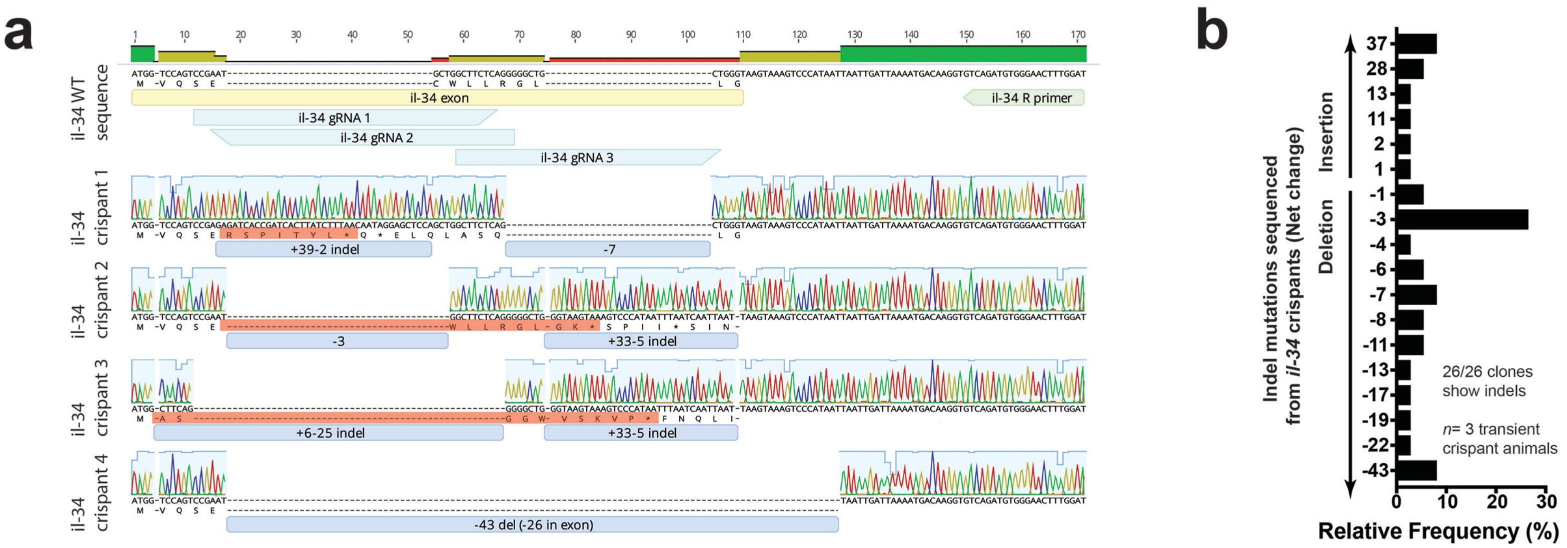

**Supplementary Figure 12.**

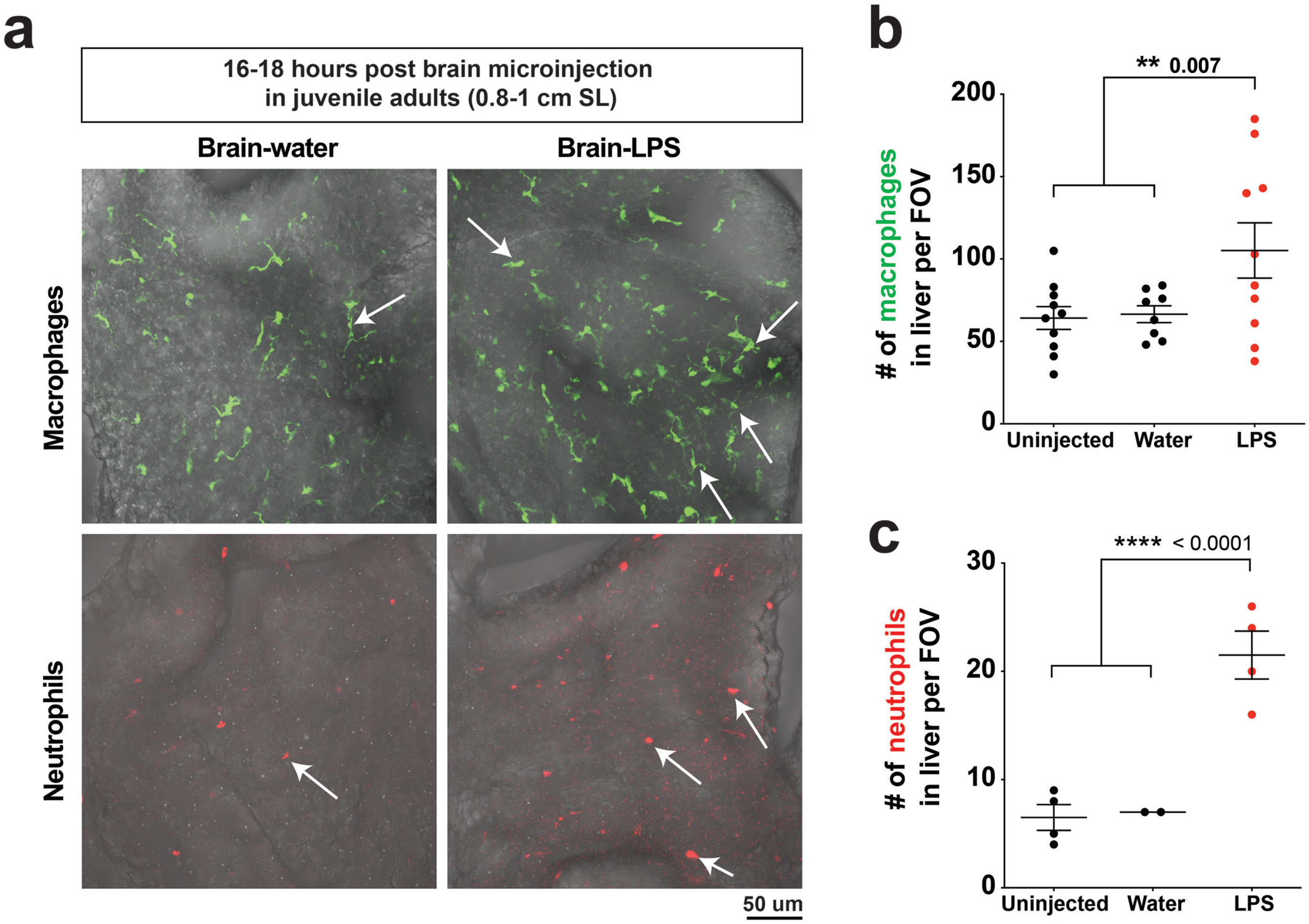

**Supplementary Figure 13.**

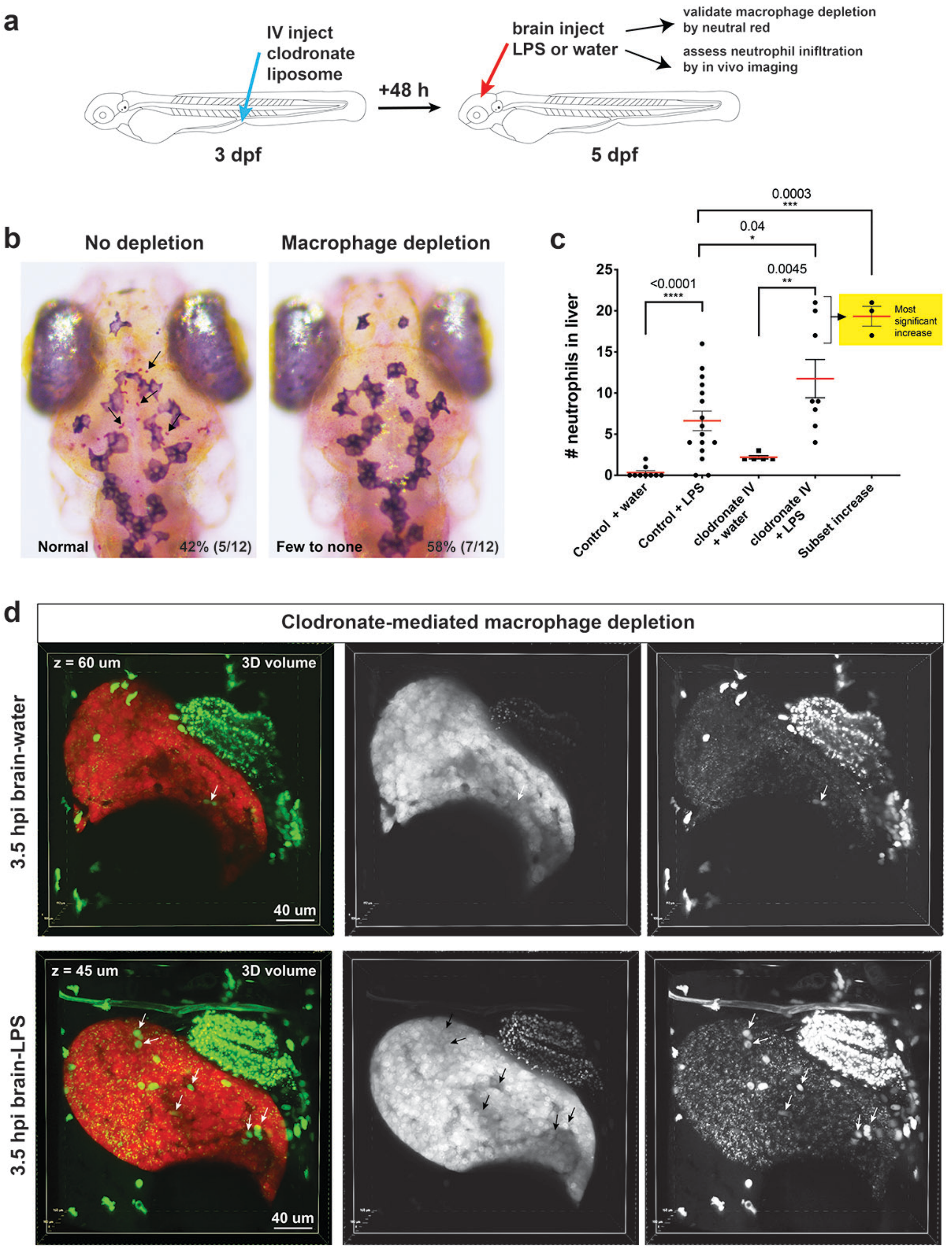

**Supplementary Figure 14.**

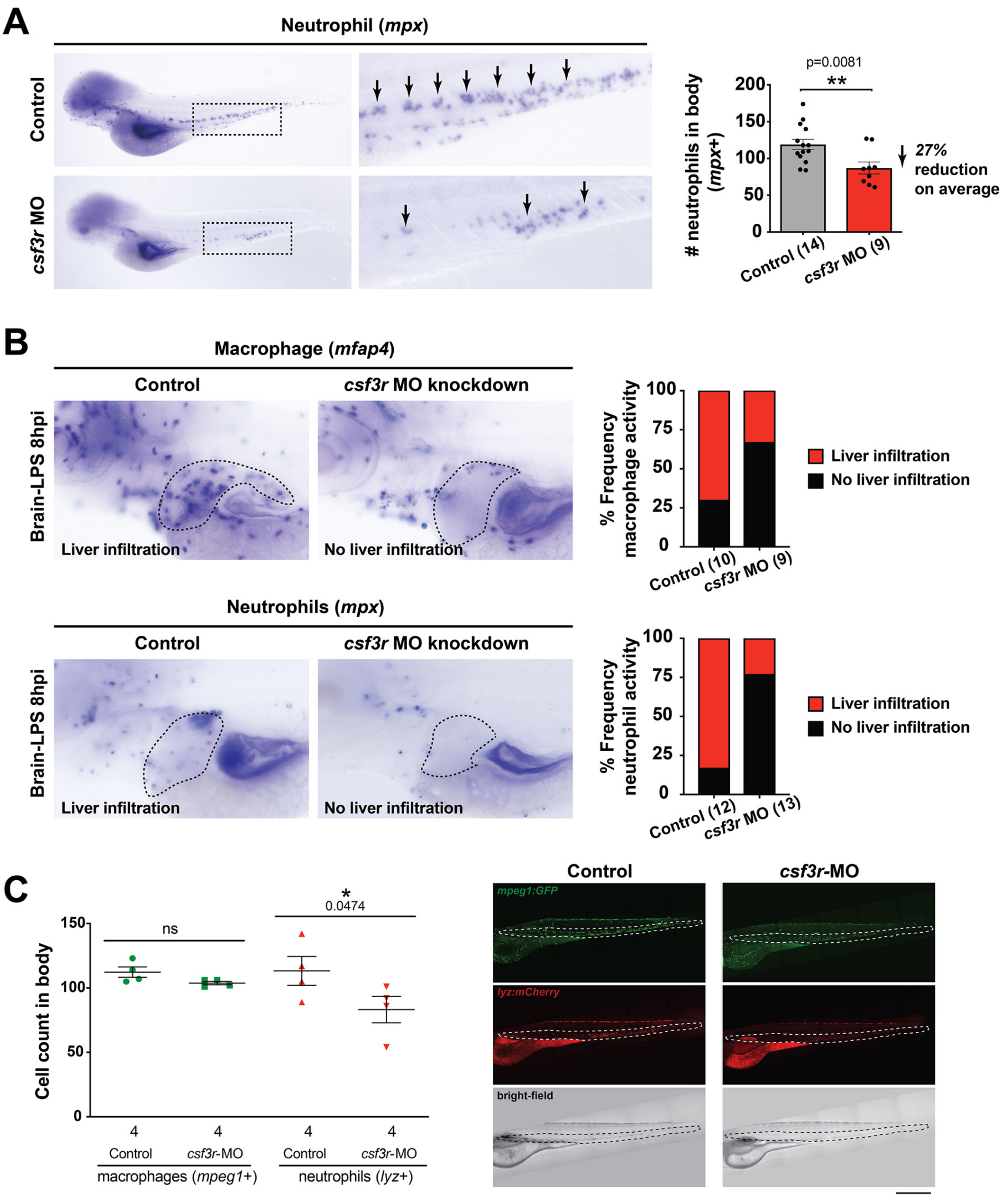

**Supplementary Figure 15.**

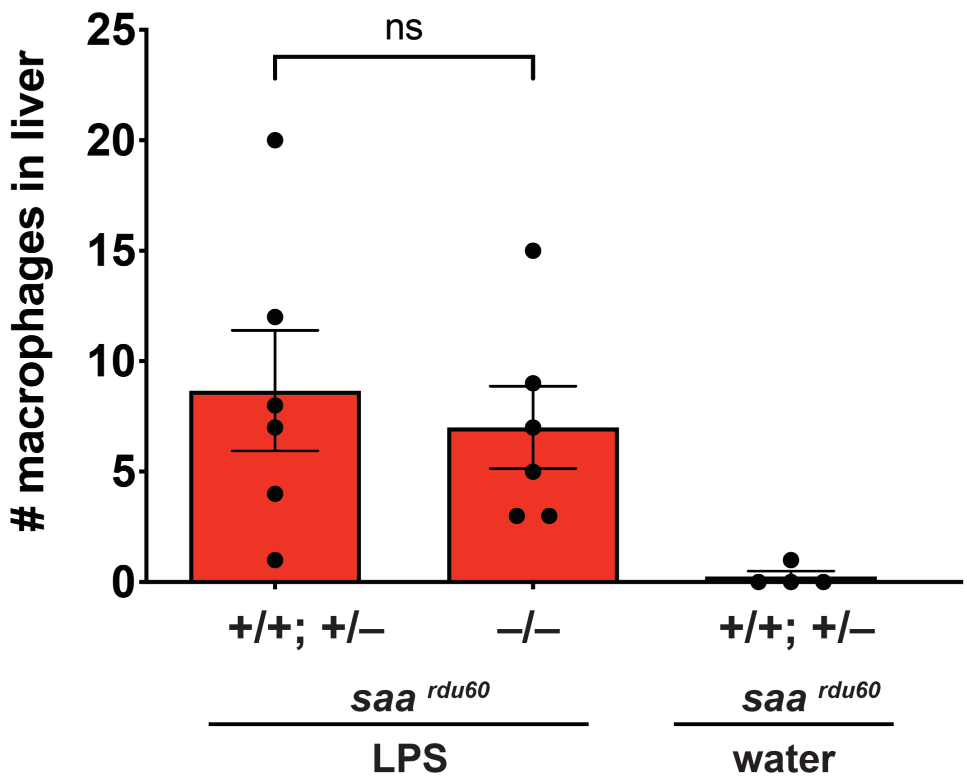

**Supplementary Figure 16.**

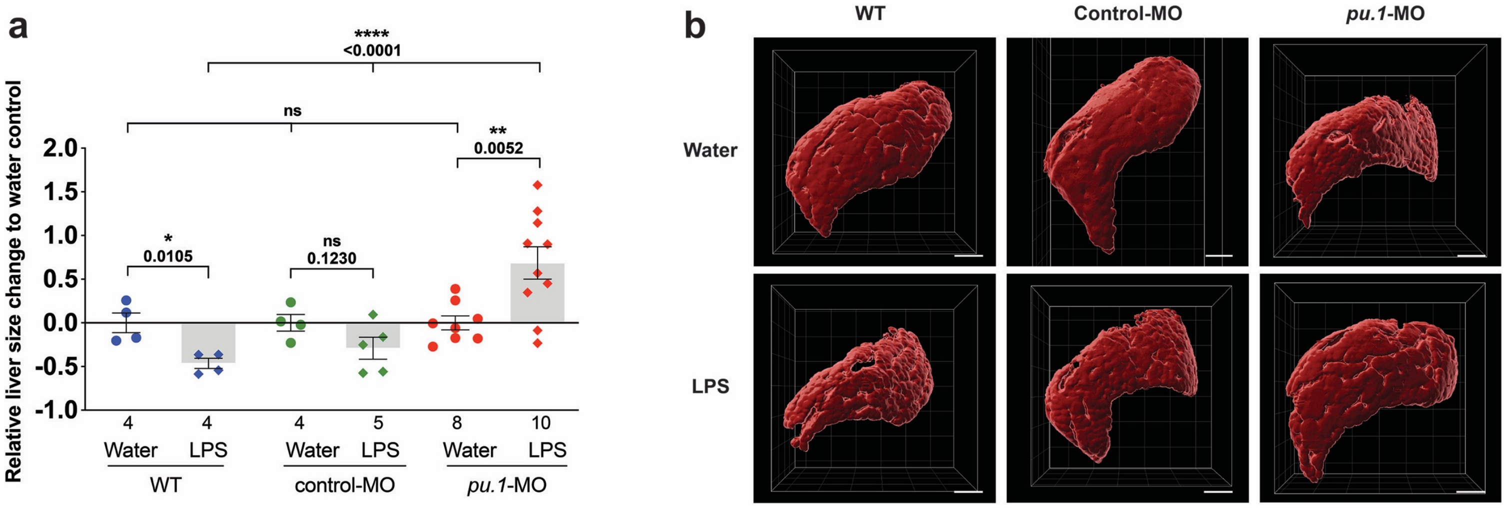

**Supplementary Table 1.**

| Primer | Sequence (5'->3') | Application | Reference |
|---|---|---|---|
| a2ml qpcr F | GGCTGTGCCATCCATAACT | qPCR | This study |
| a2ml qpcr R | CTCCGTTCCTTCTTCTGTAACC | qPCR | This study |
| ahr1a qpcr F | CTTCTGGTTTTCTTGCCGTG | qPCR | This study |
| ahr1a qpcr R | GCCATCTCCTGTACTCTCATTC | qPCR | This study |
| csf1a qpcr F | AGTGCTTCATCCTTGTCCTG | qPCR | This study |
| csf1a qpcr R | TGACAGAGTGCTTACATGGAC | qPCR | This study |
| csf1b qpcr F | TGTCCAGGCACTTCAAAGAG | qPCR | This study |
| csf1b qpcr R | GGTTAGTTGAGTTACAGTCGGG | qPCR | This study |
| ef1a qpcr F | ACATCCGTCGTGGTAATGTG | qPCR | Earley et al., 2018[2] |
| ef1a qpcr R | TGATGACCTGAGCGTTGAAG | qPCR | Earley et al., 2018[2] |
| gp130 qpcr F | GATGGAGAGGTGGAGATGATTG | qPCR | This study |
| gp130 qpcr R | TGCGAAGGCAGAGCATTATGG | qPCR | This study |
| gstp1 qpcr F | TTGAAGAGTGGATGAAGGGC | qPCR | This study |
| gstp1 qpcr R | TTTTCGACCCAGATGTCTCAG | qPCR | This study |
| il1b qpcr F | ACCGGCAGCTCCATAAAC | qPCR | Earley et al., 2018[2] |
| il1b qpcr R | GGTGTCTTTCCTGTCCATCTC | qPCR | Earley et al., 2018[2] |
| il-34 qpcr F | CCATCTGGGATCACTTGACTG | qPCR | This study |
| il-34 qpcr R | GTCATCCTCTTTTGTGAAGCAC | qPCR | This study |
| irg1 qpcr F | ACTGGGAGTGGGTTTGATTG | qPCR | Earley et al., 2018[2] |
| irg1 qpcr R | GCATACTGAGGTGGAAGAGATG | qPCR | Earley et al., 2018[2] |
| MyD88e1F | TCTTGACGGACTGGGAAACTCG | RT-PCR | Bates et al., 2007[3] |
| MyD88e5R | GATTTGTAGACGACAGGGATTAGCC | RT-PCR | Bates et al., 2007[3] |
| saa qpcr F | GCTGTGCTGGTGATGTTTATG | qPCR | This study |
| saa qpcr R | GTTCTTCCAATTGGCCTCTTTC | qPCR | This study |
| tnfa qpcr F | GCGCTTTTCTGAATCCTACG | qPCR | Earley et al., 2018[2] |
| tnfa qpcr R | TGCCCAGTCTGTCTCCTTCT | qPCR | Earley et al., 2018[2] |
| il34 genotyping F | CAGGGCATTAAGAAGGTCTTAC | genotyping (*il34^re03^*, 5bp deletion) | Kuil et al., 2019[4] |
| il34 genotyping R | CAAATGATATCATTGTTCTAAC | genotyping (*il34^re03^*, 5bp deletion) | Kuil et al., 2019[4] |
| myd88 genotyping F | GTAACGCGGAGATATACAACAAC | genotyping (*myd88^b1354^*, 4bp deletion) | Burns et al., 2017[5] |
| myd88 genotyping R | GAAGCGAACAAAGAAAAGCAA | genotyping (*myd88^b1354^*, 4bp deletion) | Burns et al., 2017[5] |

| | | | |
|---|---|---|---|
| saa genotyping F | CATGAAGCTTCTTCTTGCTGTGCTG G | genotyping (*saa*$^{rdu60}$, 22bp deletion) | Murdoch et al., 2019[6] |
| saa genotyping R | GGAATAACAACTGACCTCCAGCGGC TTCTCC | genotyping (*saa*$^{rdu60}$, 22bp deletion) | Murdoch et al., 2019[6] |

| **Morpholinos (MOs)** | **Sequence (5'->3')** | **Function** | **Reference** |
|---|---|---|---|
| csf3r/gcsfr | ATTCAAGCACATACTCACTTCCATT | Splice-blocking | Hall et al., 2012 |
| irf8 | AATGTTTCGCTTACTTTGAAAATGG | Splice-blocking | Li et al., 2011[7] |
| myd88 | GTTAAACACTGACCCTGTGGATCAT | Splice-blocking | Bates et al., 2007[3] |
| pu.1/spi1b | GATATACTGATACTCCATTGGTGGT | Translation-blocking | Rhodes et al., 2005[8] |
| tnnt2 | CATGTTTGCTCTGATCTGACACGCA | Translation-blocking | Sehnert et al., 2002[9] |
| p53 (negative control) | GCGCCATTGCTTTGCAAGAATTG | Suppresses MO-induced apoptotic effects | Robu et al., 2007[10] |

| **Small-molecule drugs** | **Final concentration used** | **Product number** | **Company** |
|---|---|---|---|
| Bay 11-7082 | 1 μM (in DMSO) | tlrl-b82 | InvivoGen |
| Celastrol | 0.22 μM (in DMSO) | ant-cls | InvivoGen |
| Dexamethasone (water-soluble) | 6.5 μM (in water) | D2915 (contains 65 mg of Dex per gram powder) | Sigma-Aldrich |
| GW2580 | 25 μM (in DMSO) *some precipitation | 72472 | Stemcell technologies |
| 17-DMAG | 5 μM (in water) | ant-dgl-5 | InvivoGen |
